# Rapid sexual reproduction in a mixotrophic dinoflagellate revealed through temporal partitioning of cellular processes

**DOI:** 10.1101/2025.04.24.649572

**Authors:** Serena Sung-Clarke, Nour Ayache, Wenguang Zhang, Mengmeng Tong, Juliette L. Smith, Michael Brosnahan

## Abstract

*Dinophysis* are specialist mixotrophs that must balance prey capture, cell division, and sexual recombination during blooms, yet relatively little is known about the occurrence and role of sex in their ecology. Here, mating was investigated in *D. acuminata* through continuous automated microscopy of cells in culture and during natural blooms. Both in culture and in situ, vegetative division and mating were phased on a diel timescale, occurring primarily at night and near dawn, regardless of prey availability. When prey were available, feeding occurred primarily during daylight hours. The sequencing of division and mating phases, the correlation of their daily amplitudes, and the timing of their establishment after a two-day light block suggests linkage of these processes. Mating, though frequent (up to 14% day^-1^ in culture), was also not associated with zygote accumulation or resting stage formation but rapid cell reproduction via meiosis. Confinement of division and mating to nighttime and early morning may minimize conflict with photosynthesis-related metabolism and/or predator exposure. Sexual reproduction was the dominant mode of proliferation during the observed blooms, accounting for 71% and 64% of new cell production in 2015 and 2021, respectively. Because it is dependent on encounter of a compatible gamete, sexual reproduction is increasingly accessible as blooms intensify. This sexual mode of proliferation may also alleviate populations’ susceptibility to pathogens, parasites, and other threats via genetic recombination, an example of Red Queen dynamics.

## 1. Introduction

Sex is fundamentally important in the life cycle of most aquatic microbial eukaryotes as their main mechanism for genetic recombination and repair. Transitions between asexual and sexual cycles reflect trade-offs between mitotic proliferation and the adaptive and physiological benefits of sexual recombination (Brawley and Johnson 1992; Azanza et al. 2018). In many algal species, sex is intertwined with physical transformations and behavioral shifts that alter susceptibility to grazing, nutrient limitation, temperature shock, and other stressors (Azanza et al. 2018). The frequency of sex and its triggers in natural populations, however, remain poorly understood. Studies of sex and its control in the ocean have primarily focused on light and nutrient-limited algal species, with more limited investigation into the reproductive ecology of prey-dependent heterotrophic microbial eukaryotes (Speijer et al. 2015; Azanza et al. 2018; Romano et al. 2022).

A basic barrier to sex is the need for cells to encounter compatible gametes. Accordingly, many species have developed aggregation behaviors (e.g., thin layer formation) that facilitate sex but also heighten resource competition, predation risk, and spread of viruses and parasites (Klausmeier and Litchman 2001; Durham and Stocker 2012). Another behavioral adaptation that can promote gamete encounter is time partitioning of sex apart from competing processes, e.g., resource acquisition and cell division (Kronfeld-Schor and Dayan 2003). In this study, time partitioning of mating is explored in the obligate mixotrophic dinoflagellate *Dinophysis acuminata*. This is one of several *Dinophysis* species that cause diarrhetic shellfish poisoning (DSP) through their production of potent phosphatase inhibitors (Ayache et al. 2023; IOC-UNESCO 2024). All *Dinophysis* species are also specialist predators of the kleptoplastic ciliate *Mesodinium*, which they retain intracellularly as their own kleptoplasts after ingestion (Park et al. 2006). *Dinophysis* can derive most, if not all, of their gross carbon requirements from these plastids during periods of prey scarcity, but cultures will not grow unless fed and exposed to light (Riisgaard and Hansen 2009; Tong et al. 2010).

Like most other dinoflagellates, *Dinophysis* are haplontic, meaning mitosis occurs in the haploid (vegetative) phase of their life cycle. Gametes are derived from vegetative cells through an unknown process and are morphologically indistinguishable (Giacobbe and Gangemi 1997). Diploid planozygotes form from fusion of paired gametes and can be differentiated from vegetative cells by their paired, rather than single, longitudinal flagella (Escalera and Reguera 2008). Cells return to the haploid vegetative stage through meiotic divisions. The proximate triggers inducing sex and return to haploid vegetative growth are not known.

In this study, the timing and extent of *D. acuminata* sexual fusions were investigated both in situ and in culture through continuous automated light microscopy using an Imaging FlowCytobot (IFCB) imaging-in-flow cytometer. Additionally, changes in cells’ DNA content during culture experiments were assessed by flow cytometry. Mating events were explored in the context of feeding and cell division timing under different prey availability and growth rates. The duration of observable division and mating stages were also parameterized so that the rates of these behaviors in blooms could be assessed from in situ IFCB records. In both field and cultured cells, vegetative division, mating, and grazing were partitioned into distinct periods each day. Notably, diploid zygotes, despite high mating frequencies, did not accumulate in culture; instead, they quickly completed two meiotic divisions, providing an alternative, rapid reproductive pathway that occurs frequently during natural blooms. These results are considered in relation to hypothesized life histories and the challenges conferred on this obligate specialist mixotroph by its reliance on *Mesodinium* prey.

## 2. Methods

### 2.1. IFCB recordings of *Dinophysis acuminata* blooms in Nauset Marsh

This study considered IFCB-derived records of *D. acuminata* blooms that occurred within the Nauset Marsh in 2015 and 2021. The marsh is made of up three, relatively deep terminal kettle ponds (6-11m) that are interconnected to each other and the Atlantic Ocean by a network of shallow (<2m) interconnected tidal channels. This area is a hot spot for harmful algal blooms in Massachusetts, USA, most prominently those of the toxic dinoflagellate *Alexandrium catenella,* which causes annual or near-annual shellfish harvesting closures (Crespo et al. 2011; Ralston et al. 2015). As a result, the smallest and northernmost of the system’s three kettle ponds, Salt Pond (Eastham, MA) has been the focus of intensive monitoring, including through deployments IFCBs since 2012 (Brosnahan et al. 2015, Fig. 1). Significant blooms of *D. acuminata* have been recorded near-annually since 2015, when IFCB observations triggered the first shellfish closure due to diarrhetic shellfish poisoning (DSP) in the northeast United States.

**Figure 1.**
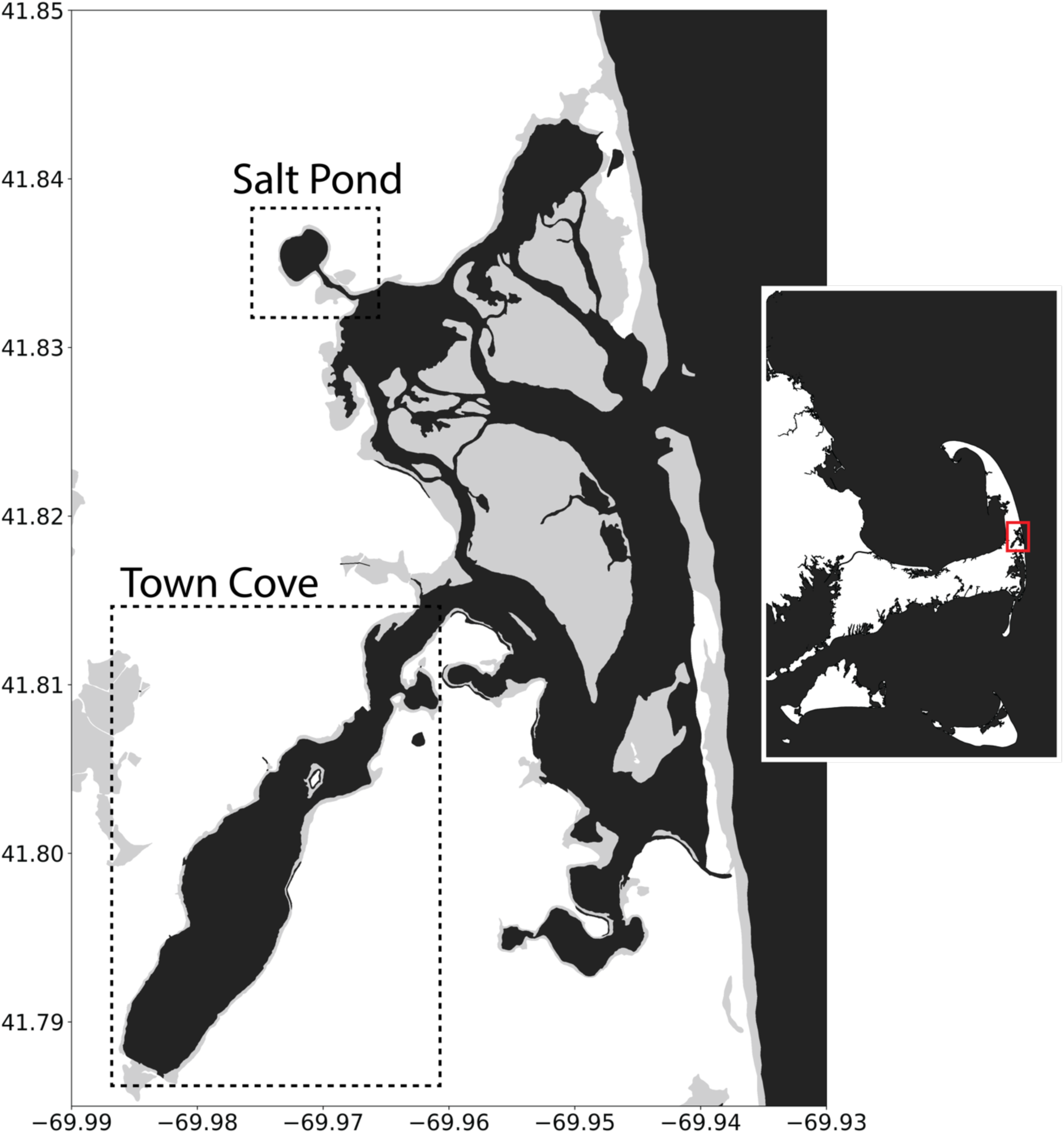
Location of Salt Pond within the Nauset Marsh system and within southeastern Massachusetts, USA (inset). Grey areas are intermittently flooded marshland.

IFCB images are sufficiently detailed to identify different *D. acuminata* cell behaviors (i.e., division, mating, feeding) as well as other microplankton and large nanoplankton (5-150 μm), which dominate Nauset’s phytoplankton communities (Fig. 2, Brosnahan et al., 2015). The frequency of division, mating, and feeding behaviors were quantified when these “events” exceeded a baseline frequency of >1 event mL^-1^ of seawater analyzed. In both years, the IFCB was deployed from a floating observatory platform positioned above the deepest part of Salt Pond and detection of *D. acuminata* was based on the cells’ red autofluorescence. During the 2015 deployment (April 10 – August 18), the IFCB was held a constant depth of ∼5 m, except for July 2-3, when it was moved along 1-m intervals between 1and 5 m depths. During the 2021 deployment (March 8 – August 2), the IFCB was moved between 2 and 3 m depths by a programmable winch. In both years, IFCBs collected samples every 20-25 minutes, with intermittent interruptions for cleaning or maintenance. Water temperature during the 2015 bloom was recorded by a HOBO logger (Onset Computer Corp., Bourne, MA) anchored near the northern shore of Salt Pond at 2.3 m depth. In 2021, temperature was measured by a profiling conductivity-temperature-depth logger that was outfitted with a phycoerythrin fluorometer (RBR Global, Inc.).

**Figure 2.**
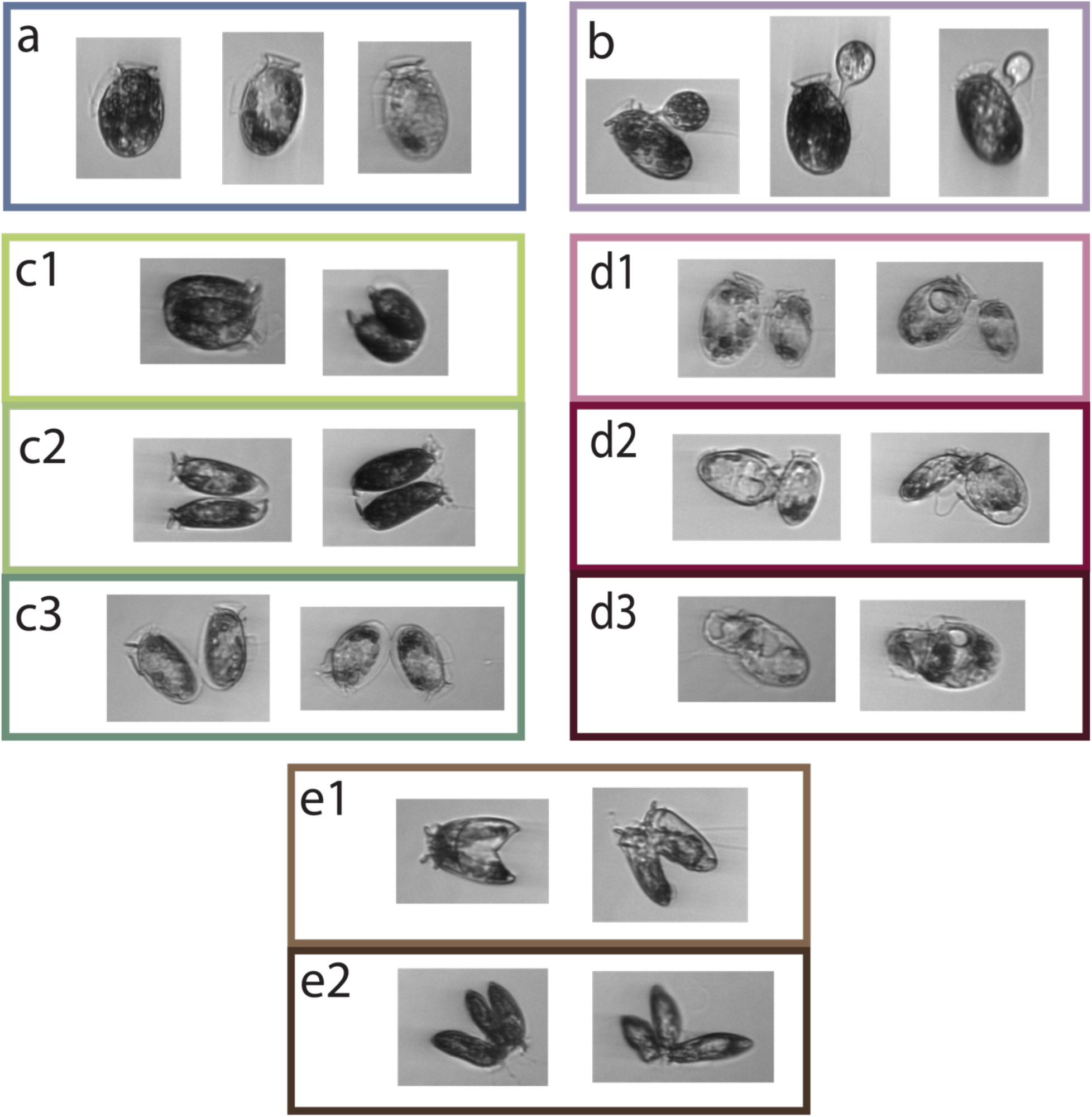
Example IFCB images of *Dinophysis acuminata* from culture: **(a)** single cells, **(b)** feeding, **(c)** dividing across the progression of **(c1)** early, **(c2)** mid, **(c3)** late division stages, **(d)** mating cells across stages of its progression **(d1)** early pairing, **(d2)** intermediate pairing, **(d3)** engulfment, and **(e)** meiotic **(e1)** dyads and **(e2)** triads.

IFCB images from a subset of all samples collected were classified through a two-step process. This subset was comprised of one sample per hour, every third day, spanning pre-bloom to post-bloom or when the time series ended (∼12% of IFCB samples collected across both years). Images were first sorted via inference with an inception_v3 convolutional neural network (CNN; Szegedy et al. 2015) model that considered 104 taxonomic groups (47 species level, 54 genus, and 3 class) and 43 additional generic, polyphyletic, and non-taxonomic categories (e.g., zooplankton, fecal pellet, fiber). Model inferences were then reviewed and corrected with a web-based annotation tool (https://ifcb-annotate.whoi.edu) for final classification of *D. acuminata* and *Mesodinium* and identification of dividing, mating, feeding, and meiotic cells (Fig. 2).

### 2.2. Culture Experiment

Temporal patterns of *D. acuminata* behavior were further explored through a culture experiment that compared frequencies of division, mating, and feeding to changes in *D. acuminata* and *Mesodinium* cell concentration at two feed ratios. This experiment also explored the role of light-dark (L:D) cycles in establishing diel phasing of these behaviors. In both experiments, *D. acuminata* strain DAMV01 (Fux et al., 2011) was maintained using *M. rubrum* strain JAMR, which in turn was fed the cryptophyte *Teleaulax amphioxeia* strain JATA (Nishitani et al. 2008a). The *Mesodinium* were grown in f/20 media without Si, a 10-fold dilution of f/2 in sterile seawater (Guillard and Ryther 1962; Guillard 1975), and fed once per week with *T. amphioxeia*. All three cultures – DAMV01, JAMR, and JATA – were exposed to 14:10 light/dark cycles under 4200k fluorescent lights (∼13 μmol photons m^-2^ s^-1^) at 15 °C. This choice was based on temperatures recorded during the Nauset Marsh blooms. Feeding of DAMV01 was stopped 4 days prior to the start of IFCB monitoring.

For IFCB monitoring, DAMV01 cells were inoculated into two 3-L jacketed beakers at an initial concentration of ∼2.5x10^5^ cells L^-1^. Cells were maintained at 15°C throughout their observation using a refrigerated circulating water bath (VWR, Model 1156D). Different prey supply ratios in the two treatments were aimed at altering their respective growth rates. The faster-growing (“prey-saturated”) culture was fed using a target ratio of 3:1 JAMR: DAMV01, while the slower-growing (“prey-limited”) culture was fed at a 1:3 target ratio. Each culture was amended with *Mesodinium* five times: once immediately preceding inoculation and then at 18, 42, 66, and 90 hours after IFCB sampling was initiated. *Mesodinium* additions were timed to occur between 7.5 and 8 hours from the start of light periods when applicable. IFCB imaging was triggered by particle side scatter. Both treatments were monitored by IFCB for a total of 12 days.

On transfer to the IFCB observation flasks, cells were deprived of light for 58 hour in an attempt to synchronize their division cycles. Following this initial light block, cells were then returned to a similar 14:10 L:D cycle as experienced prior to transfer to the flasks (30 μmol photons m^-2^ s^-1^ from 4200 K fluorescent tube lights). IFCB samples were taken every two hours during the light block, then hourly once L:D cycles resumed. To ensure homogeneity, a caged magnetic stirrer was programmed to mix the cultures for 3 minutes prior to each IFCB sample withdrawal. Sampled media was replenished daily with f/20 media, either alongside prey additions (until hour 90) or with media alone (from hour 90 through remainder of the experiment). Approximately one hour prior to each replenishment, triplicate 1-mL samples were taken and fixed with Lugol’s Solution to quantify DAMV01 and JAMR concentrations via microscopy on a gridded 1-mL Sedgewick Rafter under 100x magnification. A minimum of 400 cells were counted per sample, or the entire slide if fewer than 400 cells were loaded. Additionally, starting ∼ 6 hours into the first L:D cycle and continuing for 96 hours (four days), triplicate 12-mL samples were collected every two hours for DNA staining and flow cytometry. Culture volumes were measured both at the beginning and end of the experiment to account for evaporation.

IFCB image classification followed a similar workflow to that applied to Nauset observations. An inception_v3 CNN model was trained on 4726 human-annotated images that defined classes for *D. acuminata* grazing, dividing, and mating behaviors, as well as *Mesodinium*, cryptophyte, detritus, and bead particles. All automated CNN classifications were manually reviewed and corrected via the IFCB-annotate tool (https://ifcb-annotate.whoi.edu). Following initial classification, images of mating and vegetatively dividing *Dinophysis* cells were further sorted according to their progression through these two processes (Fig. 2).

### 2.3. Flow cytometry analysis of DNA content

Flow cytometry samples collected during the first four L:D cycles (hours 54-150) of the culture experiments were centrifuged at 2800 rcf for 5 minutes, fixed in 1% glutaraldehyde seawater solution for 45 minutes, re-pelleted (5 minutes, 2800 rcf), then stored in 1-mL of ice-cold methanol at -20°C for 4-5 days. Prior to analysis, samples were centrifuged at 4000 rcf for 5 minutes. Pellets were washed in cold filtered seawater, resuspended in a staining buffer (filtered seawater, 1% Triton-X-100, 100 μg/mL RNaseA, and 0.5 μg/mL DAPI), and incubated at 4°C overnight. DAPI fluorescence was measured using an Attune CytPix imaging flow cytometer (ThermoFisher) (405 nm laser and 440/50 bandpass filter), targeting ≥ 1000 *Dinophysis* events per sample. Flow cytometry data were gated and analyzed with the R Bioconductor tool *flowCore*. Cell cycle distributions were assigned iteratively using an algorithm based on the Watson Pragmatic Algorithm, where G1 and G2 presumed to be normally distributed, and the S curve was defined with an error function (Watson et al. 1987). Diploid planozygotes, assumed to overlap fluorescence with G2-phase cells, were grouped in the 2N distribution alongside the G2/M populations. The three distinct DNA content groups were thus initially described agnostically as N (G1 haploid cells), S (DNA replication), and 2N (G2/M or zygotes).

### 2.4. Division, Mating, and Feeding Duration Calculations

The durations of division, mating, and feeding processes were estimated through comparison of their frequencies in culture experiments to inferred rates of cell accumulation that accounted for daily media replacements and evaporative losses. Division and mating frequencies (*f_d_* and *f_m_*, respectively) were calculated by dividing the number of cells displaying each behavior by the overall count of *Dinophysis* cells per sample. Grazing frequency, *f_g_,* was similarly calculated by dividing the number *Dinophysis*-*Mesodinium* interaction images by the overall count of *Mesodinium* cells.

Net cellular accumulation, *a*, in the culture flasks was calculated as a function of the change in dilution-adjusted concentration over time:

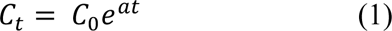

Both division and mating processes contribute to net cell accumulation. Additionally, each prey addition diluted the culture due to the added media volume. Thus, net accumulation in cell concentration is a function of the division rate (*μ_d_*), the mating rate (*μ_m_*), and the dilution rate (β):

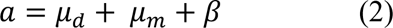

The division rate, *µ_d_*, was estimated from *f_d_* using an equation adapted from Chisholm (1981), where *t_d_* is the duration of observable vegetative cell division and *n* is the number of samples. When looking across a single day,

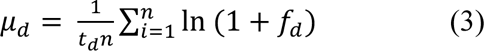

Similarly, the mating rate, *µ_m_,* is also a function of *f_m_,* its duration (*t_m_*), and number of samples observed (*n*). Here, mating was considered as both a loss and a growth process. In the former case, when two cells mate, the cell abundance is reduced by one:

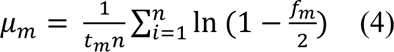

Alternatively, if mating pairs yield four haploid daughters within an observation period, mating is a source of population growth (i.e., the first step in sexual reproduction). In such cases, mating frequency (*f_m_*) and duration (*t_m_*) can then be used to calculate a sexual reproduction rate (*µ_s_*) as in Eq. 3, and this rate is then substituted for *m_m_* in Eq. 2.

The division and mating durations of *D. acuminata* were parameterized using a grid search that compared derived and observed *Dinophysis* accumulation through the first eight L:D cycles. Observed *Dinophysis* accumulation each day was calculated from Eq. 1, while derived accumulation was calculated using Eq. 2-4 across a range in durations (*t_d_* and *t_m_* each ranging from 20-180min, with 1 min intervals). Day periods were defined as starting from 10 hours after lights-on. Goodness of fit was assessed by Passing-Bablok regression and paired t-test comparisons of differences between predicted and observed changes in *D. acuminata* abundance. Parameter combinations failed the paired t-test criterion if they produced differences between predicted and observed changes that were significantly different from 0. Parameter combinations were rejected by Passing-Bablok regression if 1 and 0 fell outside the 95% confidence intervals of slope and intercept estimates, respectively.

A second method for estimating division and mating durations was based on phasing of their early and late stages. Time-of-day distributions were converted to a polar projection and smoothed by kernel density estimation (KDE) with a bandwidth determined by Silverman’s rule of thumb (Silverman, 1986). Differences in the peaks of early and late stages are estimators half the duration of the overall process. Based on an assumption that each cell or pair of cells completes the division and mating processes once begun, the relative frequencies of successive stages also reflect their relative durations (Supplementary Fig. 1).

Feeding duration was estimated from the prey-saturated experiment using observations from the last refeed on Day 4 through Day 10. The reduction rate of *Mesodinium* was first determined using Eq. 1, then this rate was then related to the frequency of *Mesodinium* being actively consumed (*f_g_*) and count (*n*) of samples to approximate the duration of observable grazing, *t_g_*, in an equation similar to that of Eq. 3 and 4.

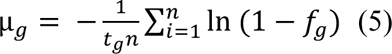

This duration, *t_g_*, is an estimate of prey handling time as applied in Type I functional response modeling (Holling 1959) and provided an additional basis to compare the observed frequencies of different *D. acuminata* behaviors.

### 2.5. Calculating proportion of cells dividing and mating

After durations of division and mating were calculated, daily rates of division and mating (*m_d_* and *m_m_* or *m_s_*, respectively) were calculated from *f_d_* and *f_m_* from both culture and in situ observations. Day periods were defined as starting 10-11 hours after sunrise (or 10 hours after lights-on). The proportion of cells undergoing behaviors resulting in growth (vegetative division with rate *m_d_* and sexual reproduction, rate *μ_s_*) were then converted to the proportion of cells undergoing these processes each day as:

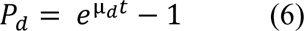

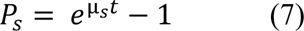

If mating is instead a loss function (with rate m_m_ < 0), the equation takes the following form:

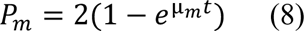

The relative contribution of vegetative and sexual reproduction was also assessed by approximating their fractional contributions to overall new cell production (*F_d_* and *F_s_*) from daily estimates of cell abundance *C_i_* and daily proportions of cells undergoing vegetative reproduction (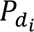) and sexual reproduction (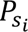).

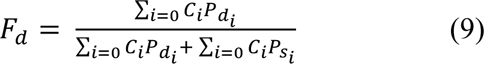

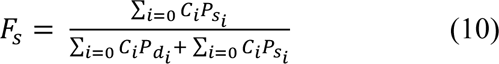

## 3. Results

### 3.1. Salt Pond bloom records

The *D. acuminata* blooms recorded in 2015 and 2021 are among the most intense and long-lasting recorded by IFCB in Salt Pond to date. The elevated *D. acuminata* concentrations observed in 2015 prompted shellfish testing by the Massachusetts Division of Marine Fisheries (MA DMF) and U.S. FDA shellfish safety programs and closure to harvesting across Nauset Marsh beginning July 15 (MA DMF 2015). Measured DST concentrations in shellfish meats rose through July, peaking at 190 ng/g OA eq. in a blue mussel sample collected on July 28 (Ayache et al. 2023). Water temperature at the HOBO logger station in 2015 (2.3 m depth) ranged from 13 to 22.5°C (mean 17.2 °C through the bloom’s development phase).

The 2021 *D. acuminata* bloom was similar to the 2015 event in seasonal timing and duration but DSTs levels in shellfish meats never exceeded the regulatory threshold for harvesting closures (Deeds and Petitpas 2021). The highest *D. acuminata* concentrations (>1.7 million cells L^-1^) were recorded by MA DMF in Town Cove. Within Salt Pond, *D. acuminata* cells were concentrated within a subsurface thin layer of elevated phycoerythrin fluorescence. This thin layer was situated below a sharp, temperature-driven pycnocline (2.2–4 m below mean sea level, averaging 3.4 m). Temperatures measured at the phycoerythrin maxima ranged from 12.2 to 17.0°C (mean: 15.1°C) through the development of the bloom. The overlaying mixed layer above the pycnocline followed a similar warming trajectory similar to the 2015 event, warming from 12.0°C to 23.2°C (mean 18.0°C) through the bloom’s development.

The IFCB record of the 2015 *Dinophysis* bloom within Salt Pond contained over 12 million images, derived from 2590 samples collected between June 1 and July 18. The highest concentration recorded by the IFCB occurred on July 3 (>1.1 million cells L^-1^) and concentrations remained >10^5^ cells L^-1^ through the end of the observational record. The volume analyzed in each IFCB sample varied from 1.0-4.2 mL (mean: 2.6 mL), adjusted based on particle concentration. 0.6% of these images were <5 µm, 66.2% were between 5-25 µm, and 33.2% were >25 µm, this largest group being the size fraction in which *Dinophysis* were observed. During the monitoring period, *Dinophysis* was detected nearly continuously (at least 21 hours daily) on 35 days. Eleven of these days were manually annotated at hourly intervals, with six days before the bloom’s peak (July 3) and five after (Fig. 3). A total of 175,206 images were annotated as *D. acuminata* cells including 923 dividing cells, 1,486 mating pairs, and 1 cell grazing on *Mesodinium*. An additional 200 images of free-swimming *Mesodinium* were recorded.

**Figure 3.**
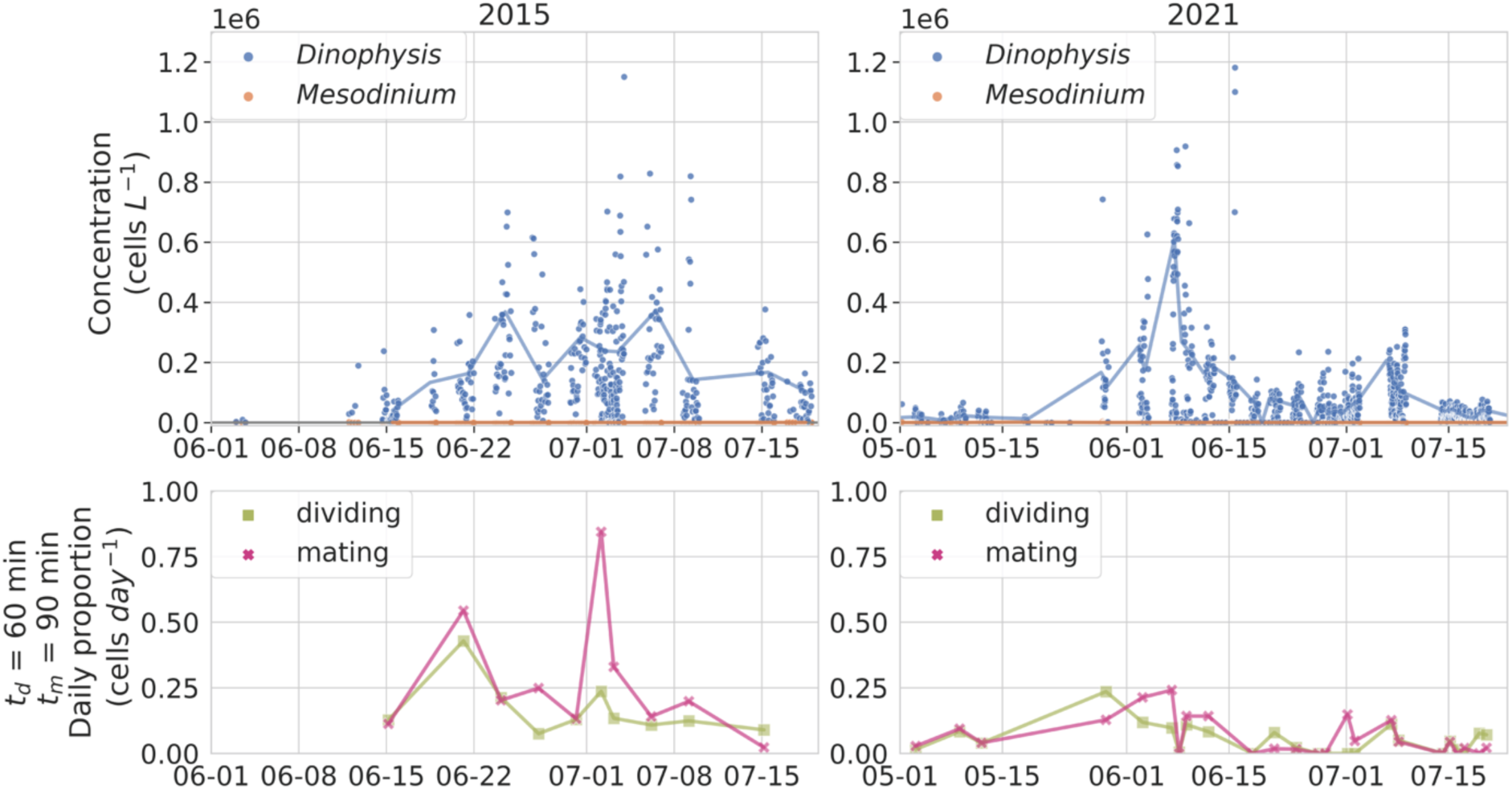
Time series of *Dinophysis acuminata* and *Mesodinium* concentrations in Salt Pond in 2015 and 2021, with estimated proportions of cells dividing or mating per day. Total concentrations were determined with hand-annotated IFCB images and known analyzed volumes. The lines represent the daily 75^th^ percentile concentration of *D. acuminata* and *Mesodinium*. Proportions are calculated from division and mating frequencies from days with full 24-hour IFCB coverage using Eq. 6 and Eq. 7.

The 2021 IFCB bloom record contained over 32 million images generated from 3259 samples collected between May 1 and July 18, 2021. The highest concentration recorded was >800,000 cells L^-1^ and occurred June 7. Daily peak concentrations then remained >100,000 cells L^-1^ through early July. The volume analyzed in each IFCB sample varied from 0.3-3.9 mL (mean: 1.2 mL). 11.2% of image particles were <5 µm, 73.1% were between 5-25 μm, and 15.7% were > 25 µm (*Dinophysis* fraction). Overall, near-continuous sampling (≥ 21 hours daily) occurred on 40 days. Samples from 22 days were manually annotated at least hourly: 5 from the bloom’s development phase and 17 from the bloom’s peak and decline (Fig. 3). *D. acuminata* cells comprised ∼ 0.5% of all images annotated (73,942 total) including 323 dividing cells, 282 mating pairs, and 1 cell grazing on *Mesodinium*. A total of 171 *Mesodinium* cells were also annotated.

During both the 2015 and 2021 *D. acuminata* blooms, cell division occurred most frequently around sunrise each day with a second, minor wave of divisions occurring shortly after sunset (Fig. 4a). Mating was similarly phased, with a single major peak occurring near dawn (Fig. 4a). Mating and division frequencies were approximately two-fold higher in 2015 compared to 2021. In both years, the highest daily mean division frequency occurred during the blooms’ development, whereas the highest daily mean mating frequency occurred at the blooms’ peaks. Daily division and mating frequencies were highly correlated in both years, with days with higher division frequency tended to also have higher subsequent mating frequency, irrespective of cell density (p < 0.001, Fig. 4b).

**Figure 4.**
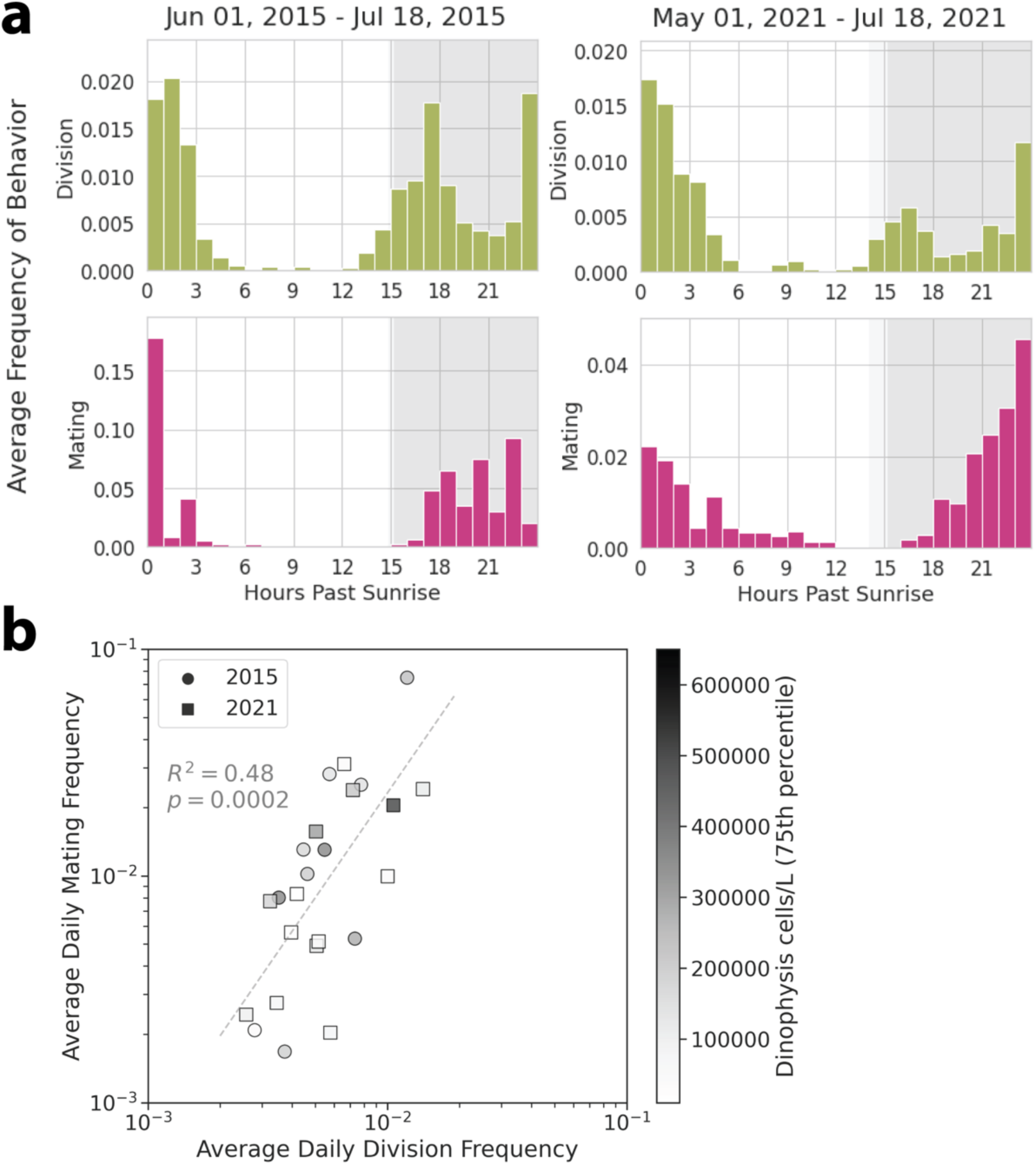
(**a**) The average frequency of division and mating behavior as a function of hours after sunrise in 2015 and 2021, defined as the count of division or mating images divided by total *Dinophysis* images. The background indicates day/night, with the light grey background indicating the earliest sunset time in that time period, and the start of the darker grey background indicating the latest sunset time in that time period. In 2015, each histogram bin represents a minimum 5 samples (mean: 14) and 551 *Dinophysis* images (mean: 5753) per hour-past-sunrise. In 2021, each bin represents a minimum 17 samples (mean: 25) and 734 *Dinophysis* images (mean: 1878). **(b)** Average daily division frequency vs average daily mating frequency in Salt Pond, for days with complete IFCB coverage in 2015 and 2021. Each day is defined as starting and ending at 2000UTC (10-11 hours past sunrise in Salt Pond). Cell density on the color scale is defined as the 75^th^ percentile concentration from all the annotated samples from that day. Linear regression was calculated with the Python package scipy.stats.

### 3.2. Culture observations

The IFCB monitored *D. acuminata* cultures under prey-saturated and prey-limited conditions through three successive conditions: (1) light deprivation with prey, (2) L:D cycling with prey, and (3) L:D cycling with progressive prey depletion. A total of 242 IFCB samples were collected from each culture condition, with analyzed volumes ranging from 1.1-1.9 mL (mean: 1.6 mL). These samples generated a total of 489,332 images. 4.7% of which were < 5 μm, 27.1% were between 5 - 25 µm, and 68.1% were > 25 µm (the *D. acuminata* containing fraction). From these, a total of 315,255 images were manually annotated: 185,311 *D. acuminata* cells (including 1423 dividing cells, 460 mating pairs, and 457 cells grazing on *Mesodinium*), 125,164 *Mesodinium*, and 8,780 detrital particles.

Across the L:D cycle periods (from hour 58 onward), the lowest daily division and mating frequencies occurred in the mid-afternoon, prompting the definition of day periods as starting and ending at this time. On day 3, several samples were excluded due to contamination in the IFCB syringe from the system’s detergent reservoir, causing elevated numbers of debris images and underestimation of *Mesodinium* (∼70-fold) and *D. acuminata* (∼2-fold) for several hours. A second gap in the IFCB record occurred between days 6 and 7 was caused by an electronic error that was resolved by power cycling the sensor.

During the initial light deprivation period (days 0-2, hours 0-58), division and mating frequencies were 0.5% and <0.2%, respectively, and lacked periodicity. Cell abundance increased only slightly, with average accumulation rates of 0.03 day^-1^ and 0.08 day^-1^ in the prey-saturated and prey-limited cultures (Fig. 5). Grazing frequency, on the other hand, was strongly phased with a 24-hour period, peaking at the end of the pre-experiment light period and shortly after prey amendments. Both prey-saturated and prey-limited cultures had similar division, mating, and grazing frequencies and similar periodicity in grazing, despite a nearly 10-fold difference in prey concentration between them.

**Figure 5.**
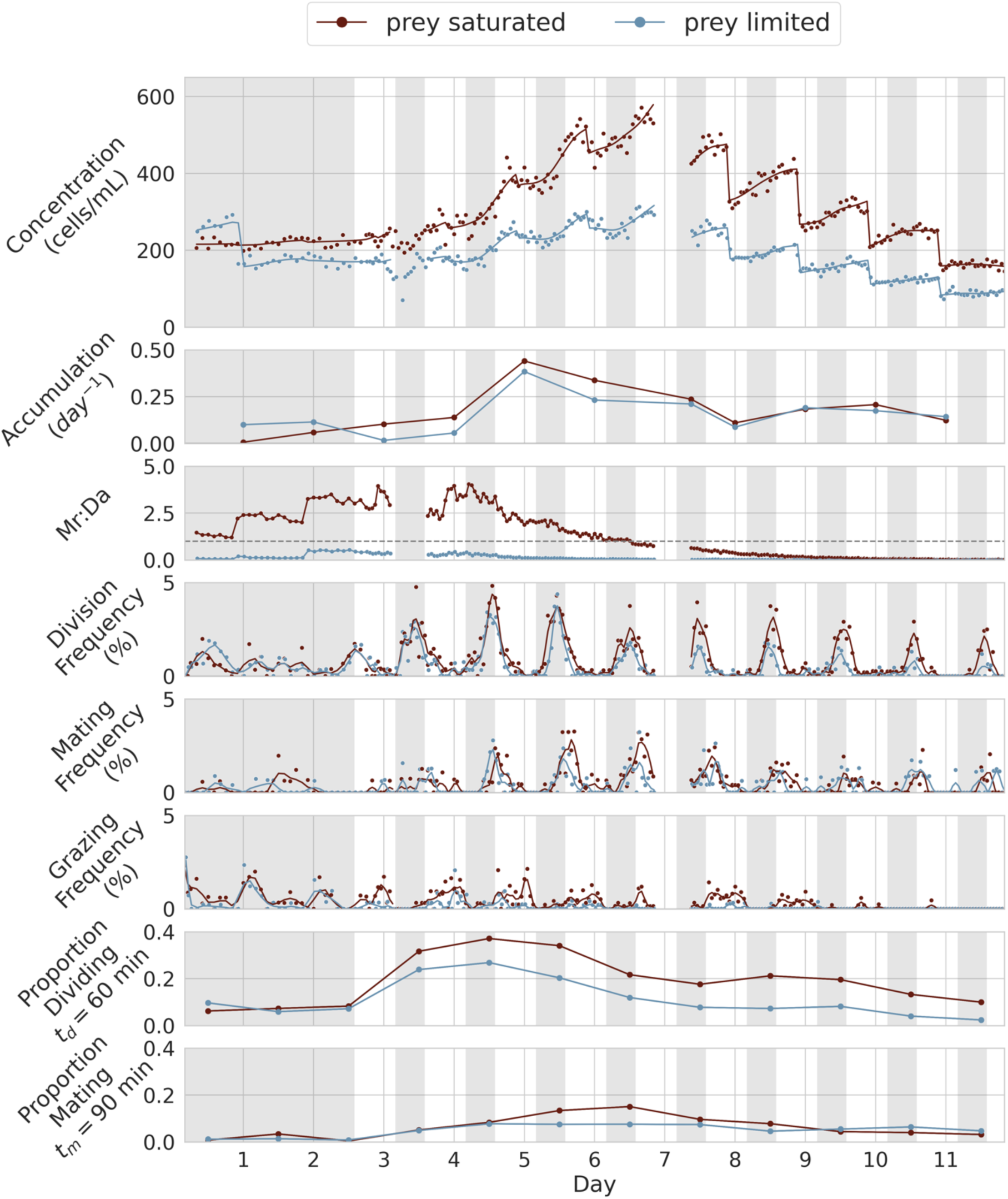
Time series of the prey-saturated and prey-limited culture treatments. Top panel shows concentration of *D. acuminata* through daily feedings and dilutions. Second panel is *D. acuminata* accumulation rates, as derived from dilution-adjusted concentrations (Supplementary Fig. 2). Third panel from top shows *Mesodinium* to *D. acuminata* ratio, where dashed line represents where the ratio was 1:1. The fourth through sixth panels are the image frequency of division, mating, and grazing behavior in *D. acuminata.* The bottom panels are the proportion of cells estimated to be dividing or mating each day (Eq. 6 and Eq. 7) given the best-constrained durations of these behaviors.

Upon reintroduction of L:D cycles, and provision of daily prey amendments (days 3–5), cells accumulation rates increased more rapidly in both prey treatments, peaking at 0.44 day^-1^ for the prey-saturated culture and at 0.38 day^-1^ in the prey-limited one (Fig. 5). This was associated with increased grazing in the prey-saturated treatment, where the average *Dinophysis* consumed ∼0.67 *Mesodinium* cells per day compared to just ∼0.1 per day in the prey-limited treatment. Phasing of division was apparent at the end of the first L:D cycle, starting around hour 74 (day 3). Mating became phased approximately one day later, with peak frequencies observed an hour into the third L:D cycle. Both division and mating doubled in average frequency during this time (1.0% and 0.4%, respectively). Grazing, which established clear periodicity earlier during the initial light-deprivation period, transitioned to a predominantly diurnal pattern (Fig. 5, Fig. 6a). Midday feedings did not set the timing of grazing, instead, it ceased at the end of each light period and resumed at the start of the next one with no prey amendment in between. *Mesodinium* were also never fully depleted during this period.

**Figure 6.**
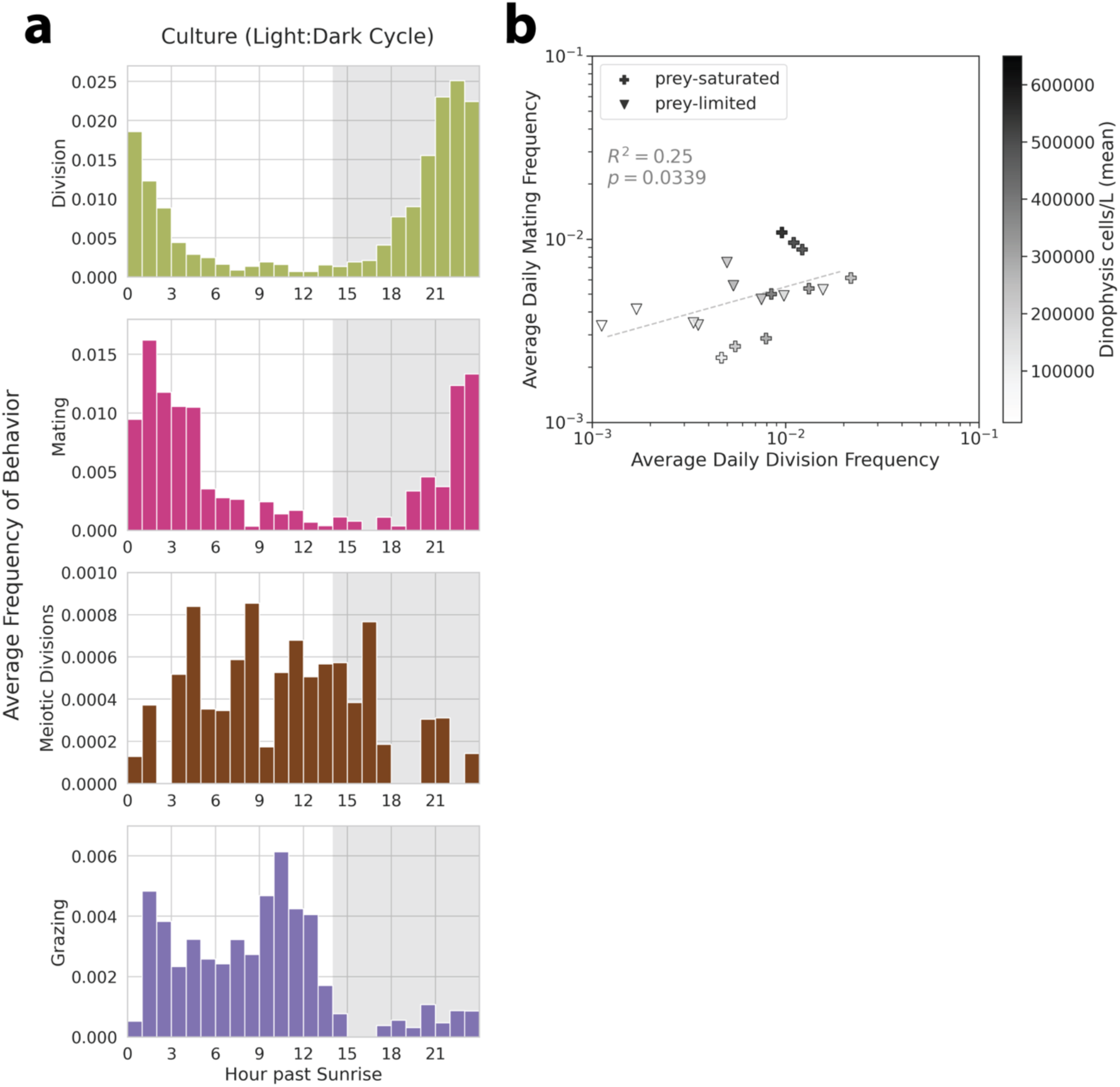
(a) The average frequency of division, mating, grazing, and meiosis as a function of hours after sunrise during the light cycle phase of the experiment, defined as the count of those observed images divided by total *D. acuminata* images. Each histogram bin represents a minimum of 14 bins (mean: 16) and a minimum 5207 *D. acuminata* images (mean: 6624). **(b)** Average daily division frequency vs average daily mating frequency in culture. Each day is defined as starting and ending at 2000UTC (10 hours past lights-on). Cell density on the color scale is defined as the mean concentration from all the annotated samples from that day. Linear regression was calculated with the Python package scipy.stats.

After prey amendments were stopped, the per-cell prey ratio declined quickly, dropping below 1 in the prey-saturated culture from day 6 until the end of the observation period. The accumulation rate in both treatments declined as well, becoming comparable starting on day 7 (0.17 day^-1^ in the prey-saturated culture and at 0.16 day^-1^ in the prey-limited culture). Despite this decline, both division and mating continued and were phased, though at lower frequencies than the earlier periods with prey and L:D cycling (averaging 0.5% and 0.4%, respectively). Division frequency was slightly higher in the prey-saturated than prey-limited treatment (0.7% vs 0.3%), while mating frequencies were more comparable (0.5% vs 0.4%). Grazing frequency also decreased but remained restricted to daylight hours.

Across all periods with L:D cycling (days 3–11), division and mating frequencies were correlated (p = 0.03, Fig. 6b). Additionally, small numbers of 3-cell ‘triads’ (8 total) and an alternative division form (‘dyads’) in which daughter cells were connected at their apical ends (81 total) were observed (Fig. 2e). These forms, similar to rare tetrads previously observed in culture (Nishitani et al. 2008b and unpubl.), were presumed to be meiotic. Unlike typical vegetative divisions or mating interactions, these forms were mainly recorded during light periods (Fig. 6a). Similar forms were observed in Salt Pond field samples, with 10 cells detected in 2015 and 1 in 2021.

### 3.3. Fate of zygotes

DNA staining and flow cytometric analysis of culture samples revealed daily cycles in the proportion of 2N cells (Fig. 7) that aligned well with IFCB-observed cell division timing. The initial light block did not fully synchronize division. Instead, the highest proportion of 2N observed was 68% and occurred 2 hours after the first dark period in the prey-saturated treatment. Daily peaks in 2N cells always preceded IFCB-observed division peaks; however, a small number of 2N cells remained in all samples, even those collected immediately after divisions had ceased. Daily reductions in 2N cells (2N_max_ - 2N_min_) were strongly correlated with IFCB division frequency (p = 0.004, Supplementary Fig. 3d). The timing of the 2N minima also followed daily peaks in mating frequency and ranged in magnitude from 3.5% to 7% of cells. Accordingly, 2N cells from daily minima were interpreted as zygotes rather than G2 or M phase vegetative cells. The amplitude of these minima did not change appreciably over the four-day time course of these measurements in spite of elevated rates of mating observed by IFCB, indicating relatively rapid reversion to haploidy by meiosis, i.e., sexual reproduction.

**Figure 7.**
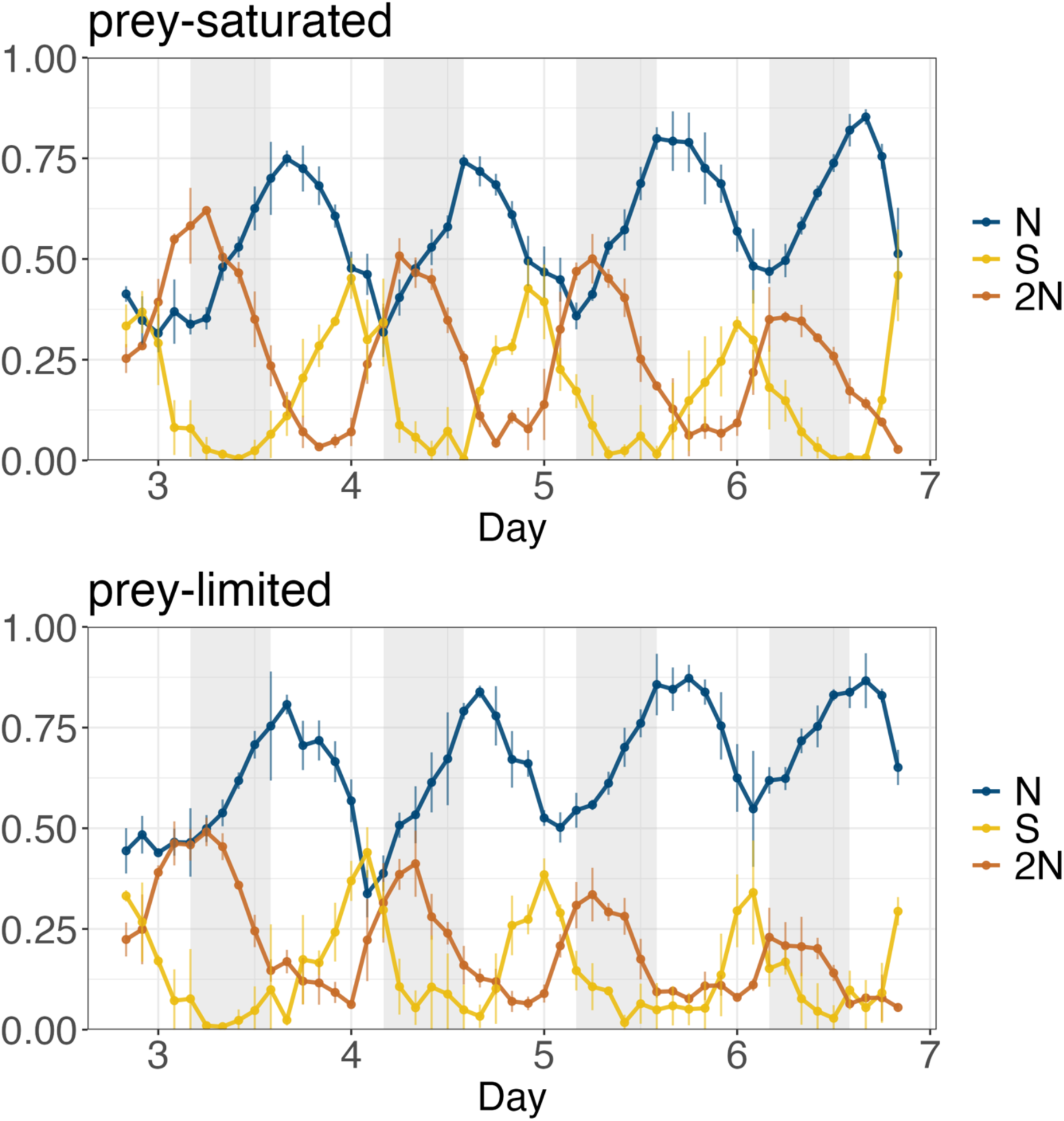
Mean proportion of cells with N (G1 haploid), S (DNA replication), and 2N (G2/M + zygote) DNA content across the first four light cycles in the prey-saturated and prey-limited culture treatments. Error bars represent the standard deviation of triplicate samples. Grey represents the dark period, and days correspond to days in Fig. 5.

### 3.4. Estimation of grazing, division, and mating durations

The observable duration of grazing by *D. acuminata* was estimated as 21 minutes from the grazing frequency and *M. rubrum* loss recorded by IFCB between day 4 and day 10 in prey-saturated culture. Similar estimation of *D. acuminata* mating and division durations was more complex because both processes affect *Dinophysis* accumulation. Further, mating can have a dual role as either a loss process (potentially halving cell concentration through zygote formation) or the initial step in sexual reproduction.

When mating was considered a loss process, the grid search constrained division duration to between 27 and 48 minutes across the full range of mating durations evaluated (20-180 minutes) and expected interdependence between mating and division duration estimates was limited. In contrast, when mating was considered a reproductive process, division and mating durations were strongly interdependent. Specifically, when mating duration exceeded 80 minutes, mitotic division was limited to a range of 50 to 90 min, in contrast, when division exceeded 80 minutes, mating was constrained to a range of 30 to 80 min (Fig. 8b). The latter result also aligned well with estimated durations developed by parsing division and mating images into early and late stages of each process.

**Figure 8.**
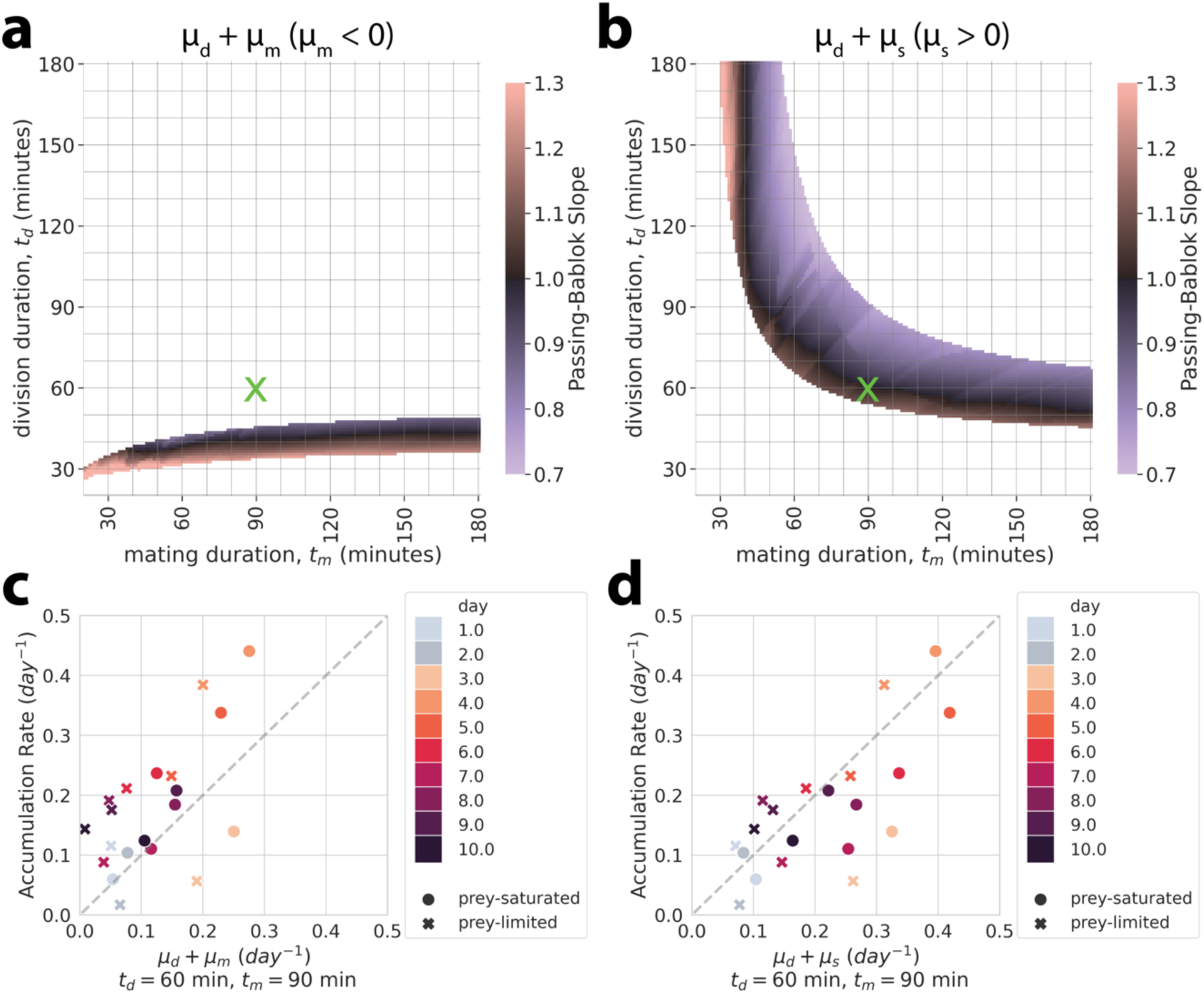
(**a-b**) Grid search fit of observed and calculated accumulation rates based on a range of division and mating durations, colored by the Passing-Bablok regression slope between those rates. (**a**) Grid search where mating is treated as a loss process (µ_m_, no subsequent meiosis). (**b**) Grid search where mating is treated as a growth process (µ_s_, meiosis follows mating). Data were only shown if observed and calculated rates were not found to be significantly different by a paired t-test (α=0.05) and if they fell within the confidence intervals (slope = 1, intercept = 0, 95% CI) for the Passing-Bablok regression. The “x” denotes the division and mating durations estimated by comparing early and late stages of these behaviors (Fig. 9) (**c-d**) Examples of one grid search comparison, using t_d_ and t_m_ of 60 minutes and 90 minutes, respectively, and comparing the calculated growth rate from these durations vs the observed accumulation rate under the assumption of mating being a (**c**) loss (µ_m_) or (**d**) gain (µ_s_) process.

In the case of mitotic division, images were split into early, mid, and late stages of division. Early-stage division images show a cleavage furrow splitting a single cell laterally (Fig. 2c1). In mid-division images, the furrow has developed to the point where daughter cells are distinct yet remain attached (Fig. 2c2). Late-division images showed nearly fully separated daughter cells both attached only at their antapical ends (Fig. 2c3). Early-stage frequency peaked earlier each day than mid- and late-stage frequencies in both cultures and field populations, consistent with expected sequencing (Fig. 8a). Greater overall frequency of the early-stage images indicated this stage is longer in duration than the latter two. Kernel smoothing analysis comparing the timing of the early- and two later stages produced a point estimate for division duration of 60 minutes (Fig. 8b).

For mating, images were likewise initially split into three stages: early-pairing, late-pairing, and engulfment. In early-pairing images, conjugated cells were attached along their dorsal axes or apicodorsal point (Fig. 2d1). In late-pairing images, the dorsal edge of the smaller cell aligned with the apical epitheca of the larger cell, with the two cells orthogonal to one another (Fig. 2d2). Lastly, engulfing cells were characterized by the larger cell subsuming the smaller one, marked by the antapical end of one gamete protruding from the apical end of the other (Fig. 2d3). The early-pairing stage images were the most frequently recorded overall, indicating that it is longer in duration than the late-pairing and engulfment stages (Fig. 8c). After lumping these later, shorter-lived stages together, mating duration was found to be approximately 90 minutes by doubling the time difference between the stage frequency peaks (Fig. 8d).

The combined point estimates from stage-timing analysis lies the near optimal fit found by grid search where mating is considered a reproductive process, but outside the parameter space that satisfied our search criteria where mating is considered a loss process. This result was consistent with the DNA staining data indicating that zygotes are short-lived intermediates of sexual reproduction. Application of the stage-timing derived mating duration indicated that 3-15% (mean 8.5%) of cells formed zygotes each day (Fig. 5), again consistent with DNA staining data (Fig. 5).

The average duration of sexual reproduction was estimated through consideration of each step in this process, starting with the formation of gametes, then conjugation and zygosis, and ending with reversion to haploidy via meiosis. Strong correlation between the daily proportion of cells dividing and mating but not cell concentration suggests that most mating cells are formed through the phased division immediately preceding their conjugation (∼5 h). Zygote lifetime was estimated as the difference between the mean daily times of mating and meiotic division. In contrast to vegetative division stages, meiotically dividing cells were rare and not tightly temporally clustered. Instead, they occurred evenly, mainly during light periods (Fig. 6a), reinforcing the conclusion that these forms are distinct from vegetative division. From assumption that daily zygote formation and meiotic division are balanced, the duration of meiotic division was estimated to be ∼18 minutes. The average time of day of these events followed peaks in mating by ∼8 h, yielding an overall estimated time required for sexual reproduction of between 12 and 13 h.

This timeframe for sexual reproduction may have been longer if gametes were rendered through prior vegetative division cycles. However, vegetative cell division was always substantially slower through the course of the culture experiments. The fastest specific rates of division observed during the culture experiment occurred on day 5 and translated to a mean vegetative cell lifetime of >2.1 days. Estimated vegetative cell lifetimes became progressively longer thereafter (>6 days by day 12). Thus, despite some uncertainty regarding gamete lifetime, sexual reproduction appears to enable a relatively faster cell production compared to vegetative division. Nevertheless, vegetative reproduction dominated culture growth, contributing 72% of new cells, while sexual reproduction accounted for 28%.

### 3.5. Contributions of vegetative and sexual reproduction to bloom development in situ

Application of the stage-timing derived duration estimates to the division and mating frequencies observed in Salt Pond indicated that large proportions of the *D. acuminata* underwent both modes of reproduction daily during the 2015 and 2021 blooms. While these proportion estimates were independent of whether mating was classified as a growth vs loss process, the high sustained mating activity throughout the bloom indicates that zygotes must have been undergoing meiosis. Therefore, in situ mating is reflective of sexual reproduction. Otherwise, the population would have been converted into zygotes several times in 2015 where mating occurred more frequently than division.

The relative contribution of vegetative and sexual reproduction shifted across bloom phases. The highest daily proportions of vegetatively dividing cells (*Pd*) were observed during bloom development phase, peaking 9-11 days prior to when highest cell concentrations were observed, nearly 43% in 2015 and approximately 24% in 2021. In contrast, mating activity was greatest during blooms peaks, with a maximum of 84% mating in a single day in 2015, and 24% in one day in 2021. As the blooms declined, the rates of both vegetative and sexual reproduction both decreased but persisted in similar proportions (average about 10% of cells per day in 2015, and 3% per day in 2021; Fig. 9). Overall, sexual reproduction contributed more to proliferation than vegetative reproduction did, representing 71% of new cell production in 2015 and 64% in 2021.

**Figure 9.**
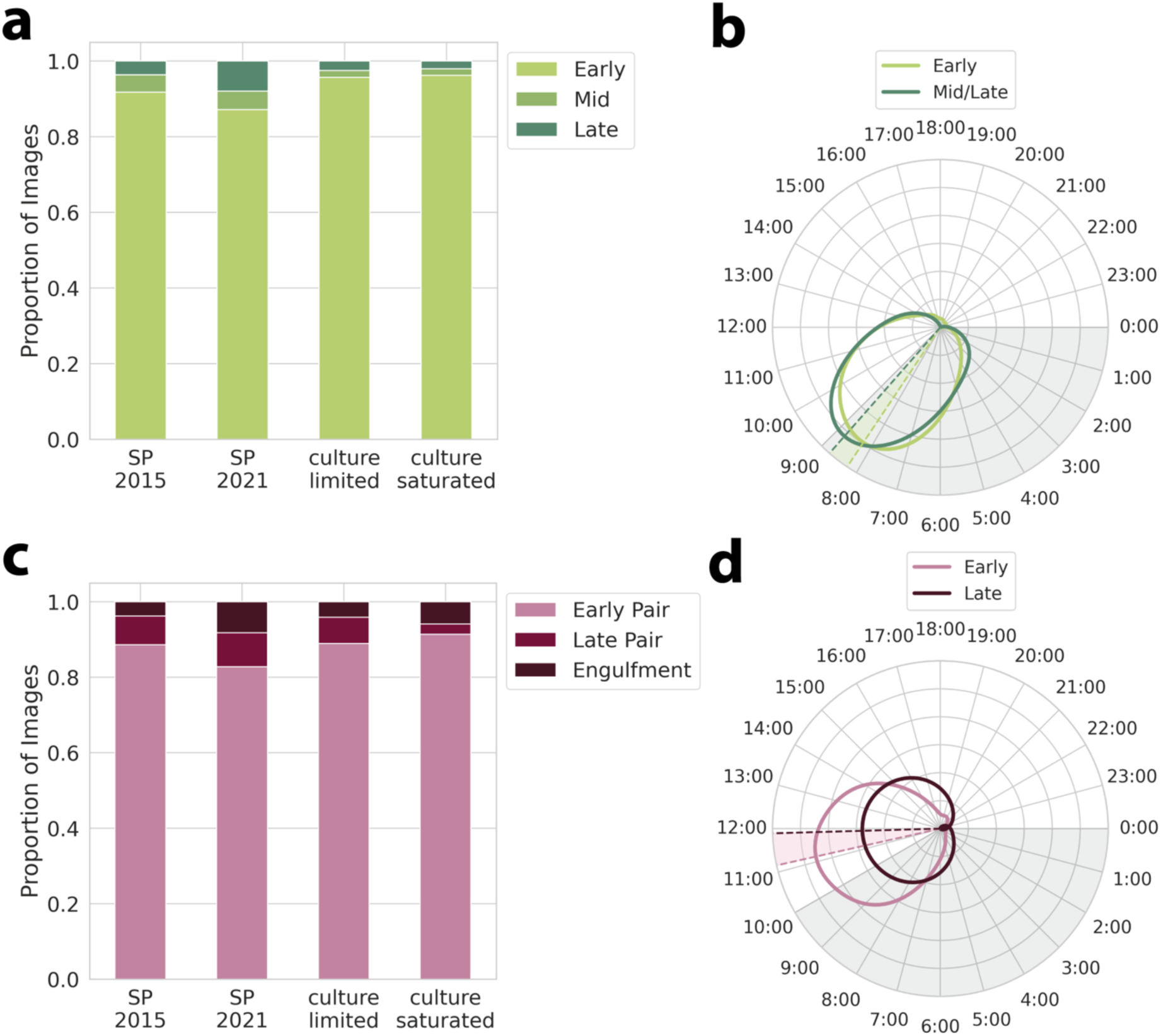
(**a**) The proportion of dividing images in early, mid, and late stages (see Fig. 2c). (**b**) Time of day distribution of early and late stage dividing cells, with a kernel density estimation (KDE) bandwidth set by Silverman’s rule of thumb. Dashed lines and shaded region show the peak observation times of early/mid and late stage dividing cells and the time difference between them (30 minutes). (**c**) The proportion of *Dinophysis* mating in early-pair, late-pair, and engulfment stages through IFCB images (see Fig. 2d). **(d)** Time of day distribution of early paired and late/engulfing mating cells. KDE bandwidth was also determined by Silverman’s rule of thumb and the shaded region shows the time difference between the time peaks in a pooled early stage (early-pair) and late (late-pair, engulfment) stages (45 minutes).

## 4. Discussion

This study aimed to characterize the occurrence, timing, and drivers of sex in blooms of the mixotroph *Dinophysis* that are nearly devoid of its obligate prey. Sexual reproduction was common throughout two blooms that produced among the highest cell concentrations of *D. acuminata* yet described (Ayache et al. 2023). While previous studies have reported mating in field populations of *Dinophysis* (MacKenzie 1992; Hajdu and Larsson 2006; Campbell et al. 2010), the sustained elevated rates of mating reported here and its linkage to new cell production highlight an underappreciated mechanism that contributes to the development and maintenance of *Dinophysis* blooms and likely many other holoplanktonic protists. High rates of mating also underscore how intense blooms like the ones in Nauset provide opportunities for extensive genetic mixing and recombination. Diel time separation of mating and division by *Dinophyis* facilitated the characterization of the rates and extent of these processes within the two natural blooms studied. Additional time partitioning of grazing and the response of these competing processes to manipulation of light cycles reveal relationships between these processes and aspects of their control.

### 4.1. Prey can promote, but are not essential, for division and sex

Consistent with prior findings (Kim et al. 2008; Riisgaard and Hansen 2009), *D. acuminata* required both light and prey to grow in culture, and cells grew faster in the prey-saturated treatment grew faster than the prey-limited one. This increased growth was supported by increased prey consumption. In contrast, *D. acuminata* in Salt Pond divided at comparable rates to cultured cells despite negligible *Mesodinium* availability. Other *Dinophysis* blooms have been observed in the absence of prey, such as the 2012 *D. ovum* bloom in Port Aransas, Texas (Harred and Campbell 2014), which is hypothesized to have been fueled by feeding prior to the bloom’s detection and observation. These Salt Pond blooms differ, however, in the persistence of their division without any significant influx of prey or evidence of feeding (Fig. 3).

Similarly, mating occurred at high rates regardless of prey availability. Comparison of culture treatments suggested prey availability could promote mating, as higher mating frequencies were observed in the prey-saturated treatment compared to the prey-limited one (Fig. 5). This finding was unexpected as many phototrophic dinoflagellates like *Gymnodinium catenatum* or *Prorocentrum cordatum* typically suppress mating when conditions favor asexual growth (Blackburn et al. 1989; Kalinina et al. 2023). It is, however, consistent with observations of the heterotrophic dinoflagellate *Polykrikos kofoidii,* which can undergo more frequent meiosis when well-fed, often bypassing its cyst stage (Tillmann and Hoppenrath 2013). The especially high and persistent rates of mating during blooms despite scarce prey as well as correlation between division and mating suggest that the effect of prey on mating rates is indirect. That is, the well-fed culture divided more because prey were available and its increased division then increased mating frequency.

### 4.2. Diel partitioning of cell behaviors as a coordinating mechanism

Temporal separation of grazing, division, and sex serves to manage interdependencies and balance opportunity costs between these behaviors. This coordination may be especially important for promoting encounters between compatible gametes. At the same time, restriction of these behaviors to certain times of day may also minimize exposure to grazers, promote more efficient prey capture, and/or manage oxidative stress associated with kleptoplast photosynthesis (Kronfeld-Schor and Dayan 2003).

Of the three behaviors assessed in the culture experiments, only grazing was periodic during the initial light block. The start of grazing periods initially coincided with prey additions, then shifted to daylight hours after introduction of L:D cycles. Two selective factors that may drive diurnal feeding by *Dinophysis* and other heterotrophic protists are competition for prey and predator avoidance (Kronfeld-Schor and Dayan 2003). If their predators are more active at night, *Dinophysis* may preferentially feed during the day to avoid them. In the heterotrophic ciliate *Strombidium arenicola*, for example, diurnal feeding can be re-established through the presence of copepodamides, a chemical signal indicating the presence of copepod mesozooplankton that are common nocturnal predators of microplankton cells (Arias et al. 2021). Copepods, being larger, typically feed in surface waters at night to avoid visual predators. Heterotrophic microplankton living in surface waters like *D. acuminata* may therefore have greater predation risk and prey competition at night.

The initial alignment of *Dinophysis* grazing with prey additions rather than the preceding L:D cycles experienced by the inoculum culture, differs from other mixotrophic and heterotrophic dinoflagellates that maintain periodic feeding for several days after switch to continuous darkness (Arias et al. 2020). The potential for prey addition to reestablish rhythmicity in the experiment’s initial phase may have been linked to starvation of experimental cells. This type of temporal control of grazing in *D. acuminata* may also be somewhat less strict, reflecting adaptation as a specialist versus their frequently scarce *Mesodinium* prey. Alternatively, prey encounters may often be correlated with light periods in situ, making prey a substitute signal for daytime. In any case, reintroduction of light cycles immediately re-established diurnal feeding, reflecting its primacy in setting and maintaining the feeding rhythm.

Daytime grazing also coincides with active photosynthesis so that carbon acquisition overwhelmingly occurred during daytime periods. Restriction of cell division and mating to nighttime may minimize conflict with carbon acquisition, yet their control showed stronger light dependence. Division and mating occurred only at low frequency and were aperiodic during the light block. The mean frequencies of both processes then increased dramatically and became phased upon introduction of L:D cycles, with phasing of division established within the first dark cycle and mating in the second. Likewise, in Salt Pond, division and mating were generally restricted to night and early morning. Similar diel patterns of division occur in the mixotrophic and heterotrophic dinoflagellates *Karlodinium armiger, Gyrodinium armiger,* and *Oxyrrhis marina,* as well as in the ciliate *Strombidium sp.* (Arias et al. 2020). However, these have different light-dependence than *D. acuminata:* division phasing in *K. armiger* persisted even in continuous darkness, while for the other species, division phasing initially ceased on switch to continuous darkness then was reestablished after about 36-48 hours. Here, *D. acuminata* division phasing was disrupted and did not reestablish itself during the 58-h dark period.

The presence of low frequency, aperiodic division during the light block indicates a light-mediated, though not strictly light-*dependent*, checkpoint in division, comparable to some photosynthetic species where procession of cell cycles is dependent on chloroplast division (Sumiya et al. 2016). *Dinophysis* differ from these in the instability of their kleptoplasts and requirement to graze to maintain light harvesting (Rusterholz et al. 2017; Garric et al. 2024). This contrasts with many algal species where light is a strict requirement for division. A prime example is the dinoflagellate *Alexandrium catenella*, which achieves division phasing through two light-dependent cell cycle checkpoints. In this species, light block followed by L:D cycling causes half the population to divide during the first L:D cycle, then full synchronization occurring on the second (Taroncher-Oldenburg et al., 1997). No similar synchronization of divisions was observed here (fewer than one-third of cells divided during the first L:D cycle) nor in previous work that applied the same light block procedure to *D*. *acuminata* (Jia et al. 2019). This discrepancy may arise from a partial checkpoint mechanism that allows more flexibility to manage prey or light limitation.

Phased nighttime division is common among phototrophic protists and may serve to optimize photosynthetic efficiency or enable more efficient cell cycling (Nelson and Brand 1979; Chisholm and Brand 1981). However, there are also many notable counterexamples, such as fast-growing diatoms and the mixotrophic dinoflagellate *K. armiger*, which divide during the day (Chisholm et al. 1980; Arias et al. 2020). For these species, daytime mitosis may offer protection from UV and/or oxidative stress during prior genomic replication or offer advantages for nutrient uptake (Nelson and Brand 1979). This diversity of strategies highlights how marine organisms balance multiple selective pressure (e.g. photosynthesis, grazing, competition, and avoidance of physico-chemical stress). In the case of *Dinophysis*, daylight is essential for photosynthesis and serves as a cue for it to feed. Why and how *D. acuminata* restricts its vegetative divisions to night periods remains to be described. Restriction of division to nighttime is not strict as meiotic divisions generally occurred during daytime periods of the culture experiment. Mating, the initial step in sexual reproduction, was most frequent near dawn and was completed in the first hours of light periods. This timing appears to be set by linkage of mating and vegetative division.

Division and mating linkage was revealed by the sequencing of their phases, the correlation of their daily frequencies, and the lag in establishment of phased mating after resumption of L:D cycling in the culture experiment. Given their linkage, mating is likely promoted by and dependent upon recent vegetative cell division (Fig. 4b, Fig. 6b). Another corollary is that the gametic phase is short-lived, most likely lasting between 4 and 5 hours (Fig. 6). Less clear is the fate of gametic cells that fail to conjugate. It may be that all newly divided vegetative cells are gametic and revert to the vegetative reproductive cycle if no compatible cells are encountered. The tendency toward gametic behavior and/or differentiation into different mating types may also depend on the internal state of newly divided cells. Most conjugated cells pairs were anisogamous (i.e., one large and one small gamete) as also seen in clonal cultures of *D. caudata* (Nishitani et al. 2008a). Size differentiation could arise through prey limitation where only some cells successfully capture *Mesodinium* or are able to meet their energy needs via photosynthesis for vegetative reproduction. Differences in the strength of division and mating linkage between the culture experiments and Salt Pond blooms lend some support for this hypothesis. In Salt Pond, where *Mesodinium* were extremely scarce in both years, linkage between mating and division was strongest as evidenced by lower division:mating ratios and tighter correlation between their daily frequencies (R^2^=0.48 vs. 0.25; Figs. 3–6). This suggests that sexual reproduction may be increasingly favored as *Dinophysis* are starved.

### 4.3. The role of sexual reproduction in bloom development

*Dinophysis* zygotes were remarkably short-lived in the culture treatments (∼8 hours). This could be determined because meiotic division forms were distinct from vegetative ones and were observed in the absence of cumulative conversion of the culture to 2N and greater DNA content. The lack of zygote accumulation through repeated cycles of mating is akin to observations from a culture of the dinoflagellate *Proroceratium reticulatum*, where a small, persistent subpopulation of biflagellated 2N cells were interpreted as cyclic generations of short-lived zygotes (Salgado et al. 2017). In this study, neither biflagellate nor a 4N cell subpopulation (indicative of meiotic chromosome replication) were recorded. The former is due to the inability to see clear flagella from IFCB imagery. The latter is likely due to the temporal dispersion of meiotic divisions over ∼12 hours each day and the relatively small proportion of zygotes overall (<10%); These factors make 4N detection difficult, especially if meiotic replication is followed quickly by formation and breakup of tetrads.

Consistent with tetrads being highly ephemeral, only derivative triad and dyad meiotic daughters were recorded by IFCB, similar to those reported in *Symbiodiniaceae* dinoflagellates (Figueroa et al. 2021). It is also possible that failure to detect tetrad forms reflects disaggregation of clusters between Meiosis I and II or sensitivity to fluid handling by the IFCB sensor. Even lower observation of meiotic forms in Salt Pond might be caused by uneven depth distribution of these forms relative to others. In *A. catenella* blooms at the same site, zygotes migrate to the surface upon formation, promoting their dispersal out of the pond (Brosnahan et al. 2017). If *Dinophysis* zygotes have similar vertical swimming behavior, they would have been undersampled by the IFCB, which collected nearly all samples between 2-5 m depths. Sexual reproduction must still have occurred within the Salt Pond blooms as without meiotic division the population could not have sustained such high frequencies of mating for weeks.

The successive processes of mating (syngamy) and meiosis comprise sexual reproduction, whereby pairs of parent gametes produce four daughter cells, providing an alternative pathway for bloom development. The sexual cycle completed in as little as 12-13 h from the formation of a gamete to production of daughter cells. In contrast, vegetative cell generation times ranged from 2.1 to >6 d (Fig. 10). Application of division and mating durations derived here showed that sexual reproduction contributed a significant portion of new cell production in the culture experiment and was the dominant source of new cells during the 2015 and 2021 Salt Pond blooms. These findings are well aligned with the “meiosis for proliferation” hypothesis from Lin et al. (2022), who recorded similar repeated cycles of meiosis during bloom development by the dinoflagellates *Prorocentrum shikokuense* and *Karenia mikimotoi*. A key facilitator of this mechanism may be the aggregation of *D. acuminata* cells in thin layers, making gamete pairing more easily accessible. However, elevated cell concentration, while pre-requisite for efficient gamete pairing, was not sufficient by itself to drive a complete switch toward sexual reproduction. Mating frequencies were variable between the culture study, the 2015 bloom, and 2021 bloom, despite comparable densities.

**Figure 10.**
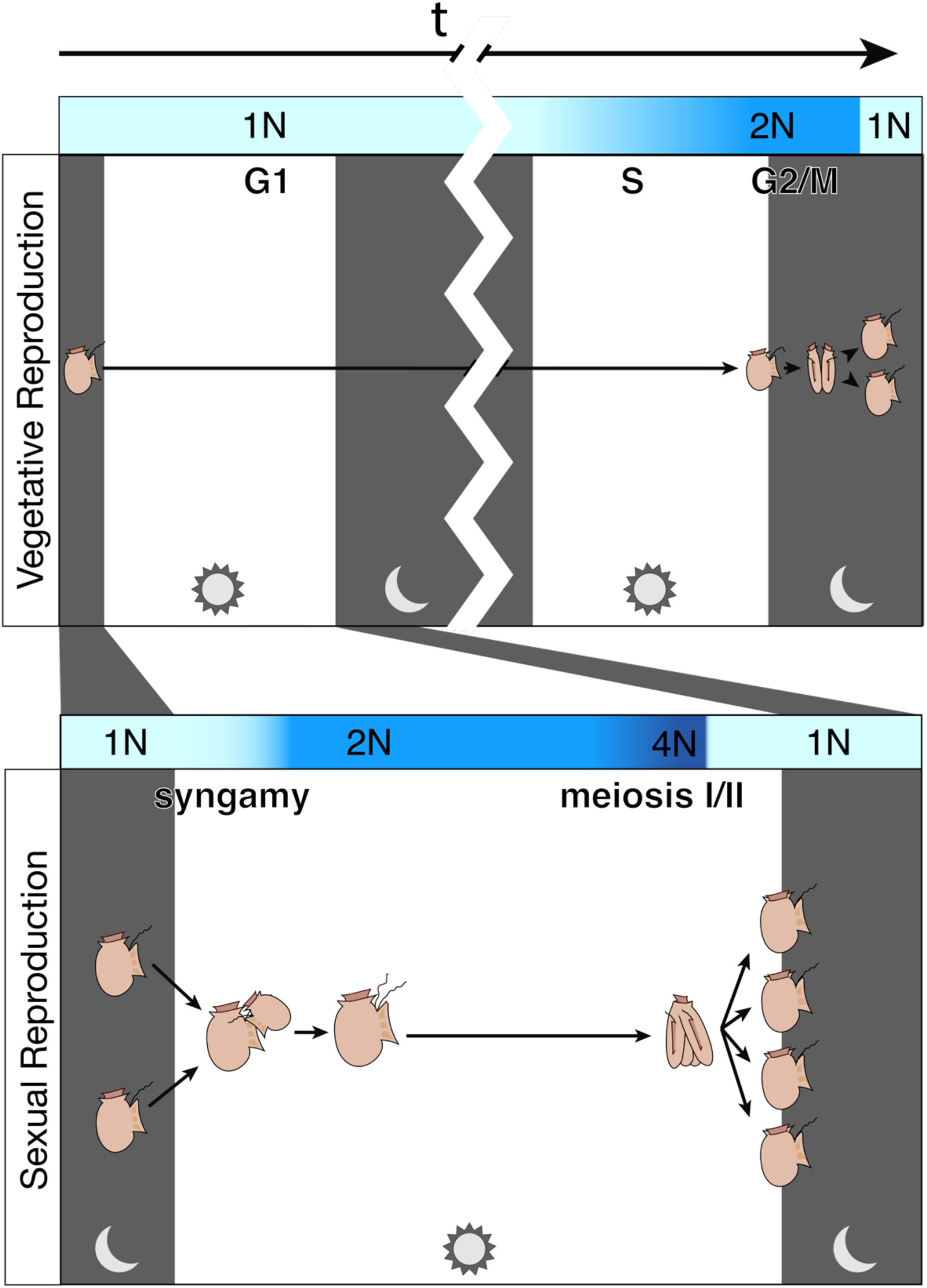
Diagram illustrating how sexual reproduction can bypass the vegetative cell cycle. “New” cells from vegetative division are generated at night. The top panel shows the progression of vegetative replication from the culture experiments here, where doubling time was >2 days, DNA replication (S phase) occurred during daylight hours, and mitosis happened at night. The bottom panel illustrates how gametes can fuse in the early morning, leading to a short-lived zygotes that undergoes meiosis later in the same day, resulting in a much quicker cell doubling.

Essential to the discovery of sex as the dominant mode of reproduction in the Salt Pond blooms was estimation of mating and division rates from IFCB records. These calculations rely on our estimates of division and mating duration, which were developed in two ways. First, expressions derived from Chisholm (1981) (Eq. 3, Eq. 4) were applied to investigate plausible ranges of division and mating duration and compare derived growth estimates to observed accumulation. While this approach successfully constrained the relationship between mating and division duration (Fig. 8), it did not yield single point estimates that could be applied to the in situ records. For this reason, a second, novel method, was developed to estimate these durations based on the timing of their distinct daily phases in the culture experiments (Fig. 9, Supplementary Fig. 1). This approach was enabled by the phased nature of these processes and the hundreds of images that captured the progression of each during the culture experiment and generated single estimates of mating and division duration that fell neatly within the bounds set by the grid search approach.

### 4.4. Sex can promote bloom resilience

Sex is ubiquitous in eukaryotic life and is as ancient as eukaryotes themselves. Very few asexual lineages exist and those examined have resulted from loss of ancient sexuality (Flot et al. 2013; Speijer et al. 2015). However, there is little consensus for why sex has persisted and remains so pervasive, especially in microbial species that may switch facultatively between asexual and sexual modes. While sex enables genetic repair and recombination, it is energy intensive, can heighten predation risk (Zuk and Kolluru 1998), and introduces the intrinsic evolutionary hazard of disrupting gene combinations that confer fitness advantages (Otto 2009). Therein lies a paradox: given its costs, why do eukaryotes undergo sex at all?

An often cited factor that ensures the persistence of sex in many protists is its coupling with behavioral and physiological transitions that make reversion to asexuality evolutionarily disadvantageous or costly (Sagan and Margulis 1987). Examples include meroplanktonic dinoflagellates, e.g., *Alexandrium*, *Pyrodinium*, and *Proroceratium* species, that couple sex to the formation of benthic resting stages for survival through periods of unfavorable environmental conditions (Salgado et al. 2017; Brosnahan et al. 2020); diatoms that rely on sex to escape a diminution trap related to the regeneration of their silica frustules (Chepurnov et al. 2004); and the coccolithophore *Gephyrocapsa huxleyi* whose haploid and diploid forms differ in their motility, growth potential, and resistance to viral infection (Frada et al. 2008). However, it’s unlikely that similar physiological coupling is responsible for maintenance of sex in *Dinophysis*. No resting cysts were produced by *Dinophysis* in this study nor observed by prior investigations of its life history (Hajdu and Larsson 2006; Escalera and Reguera 2008; Reguera et al. 2024). *Dinophysis* also do not rely on sex to avoid depauperating, as small cells can grow without being hindered by inherited frustules or thecae (Reguera and González-Gil 2001). Further, the idea that asexual and sexual stages may provide differing selective advantages is undermined by the short lifetime of the *D. acuminata* zygotes and similarity of its diploid and haploid forms. Rather, sex likely persists because it confers advantages directly rather than via a byproduct of another process.

It’s noteworthy that mating rates reported here were all associated with exceptionally high cell concentrations (>10^5^ cells L^-1^). For comparison, more typical maximum densities observed during blooms are 1-2 orders of magnitude lower (Ayache et al. 2023). These high concentrations both facilitate sex and enhance the value of the genotypic diversity. According to Red Queen dynamics, organisms are constantly in an arms race with biological antagonists (e.g., parasite, predators, competitors). Genetic variation promotes adaptability that is required to maintain relative fitness (Van Valen 1973). Whereas a population of genetically similar individuals generated through vegetative growth and selection will become increasingly prone to pathogen species, one that grows through sexual reproduction will maximize its genotypic diversity and thereby hinder selection and adaptation of its antagonists (Lively 2010). Among these are viruses, parasites, and bacteria whose potential for proliferation within a *Dinophysis* bloom is enhanced by the bloom’s density. Sex then provides a mechanism both for disrupting the success of these pathogens while also continuing to multiply (Hamilton 1980).

Prioritization of sex by individual *Dinophysis* cells appears to be modulated by changing costs and benefits of sex relative to division. In early bloom development, when prey competition is low, specialized pathogens scarce, and compatible gametes relatively rare, they favor vegetative division. Later, as concentrations increase, sexual reproduction becomes accessible as a fast way to reproduce while also generating new genotypic diversity (Fig. 11). The transition to sexual reproduction could rely in part on extrinsic cues, such as high density of conspecifics or pathogen threat. When planozygotes of the cyst-forming dinoflagellate *Alexandrium minutum*, for example, are infected by the parasite *Paryilucifera sinerae*, they tend to rapidly undergo meiosis rather than forming cysts, thereby accelerating recombinant offspring production and enhancing resistance to infection (Figueroa et al. 2010). Associated recombination not only guards against antagonistic interactions with other organisms but also helps populations contend with changing physico-chemical conditions.

**Figure 11.**
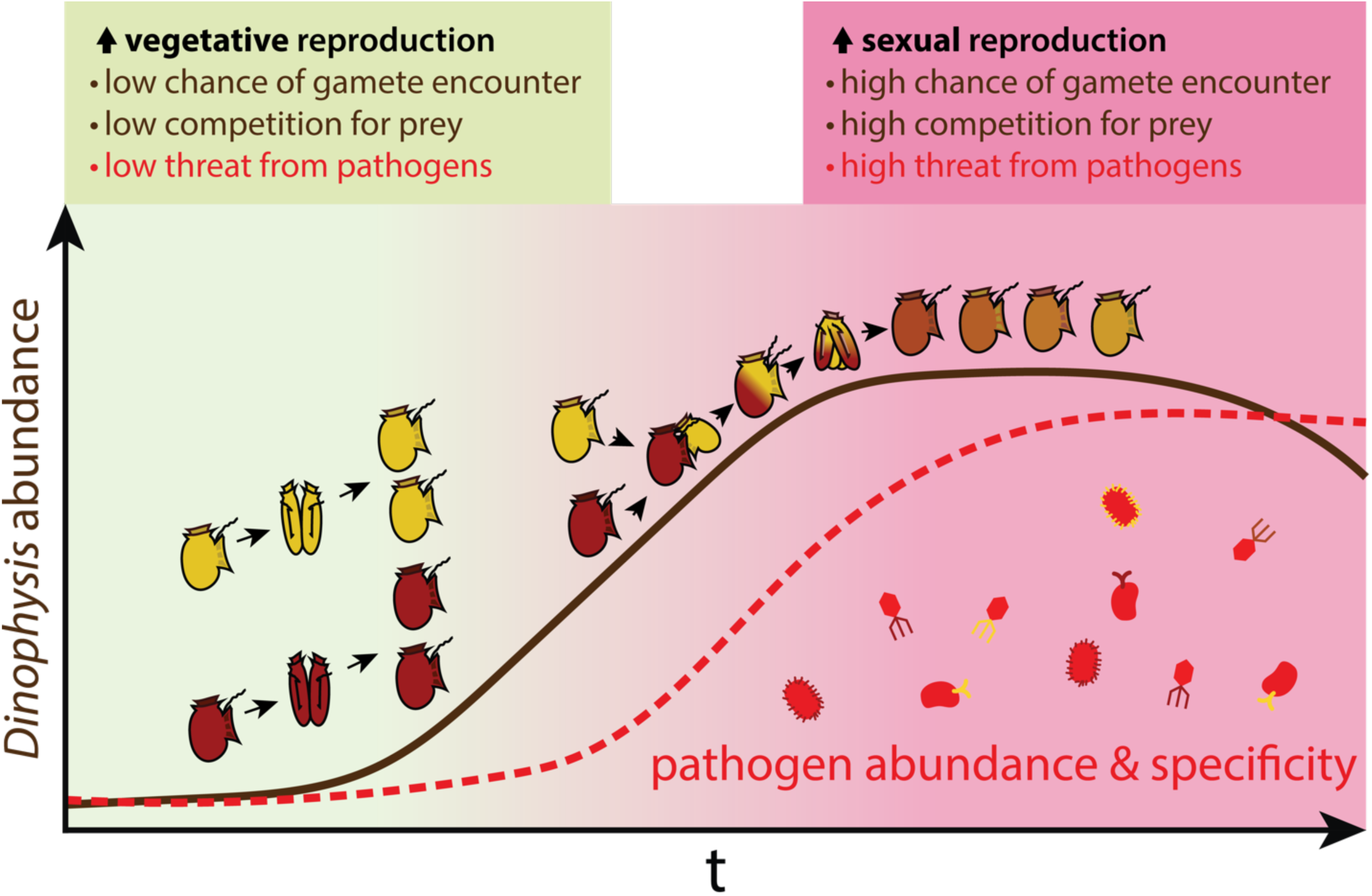
Illustration of a shift from vegetative to sexual reproduction during a bloom, an example of Red Queen dynamics. In early stages of a bloom, cells are more likely to undergo vegetative reproduction due to low gamete encounter, prey competition, and threat from specific pathogens. As the population grows, prey are depleted and gamete encounter increases; additionally, short-generation pathogens may become increasingly targeted to abundant, asexually generated genotypes. Sexual reproduction leads to genotypically diverse offspring, some of which are then more likely to elude infection.

Sex, therefore, is not only an opportunistic, but protective, behavior during *Dinophysis* blooms, contributing to a population more adaptable to biotic threats and environmental changes. For *Dinophysis* and other organisms to realize these potential benefits of sex, they must maintain behaviors that maximize their success in encountering and coupling with compatible cells. The nighttime timing of mating is one such adaptation. To the extent that either timing of gametogenesis or syngamy vary, barriers to genetic recombination can develop, isolating sub-populations, and potentially driving sympatric speciation. Conservation of the timing of sex and/or its linkage to the timing of phased divisions, may be essential for *D. acuminata* to adapt to changing conditions and threats during blooms.

More broadly, *Dinophysis* are one example of many marine protists that aggregate and bloom within coastal features through the interaction of their behavior, external physical forcing (e.g., tidal trapping, fronts, etc.), and nutrient enrichment. These coastal habitats therefore provide opportunities for cells to sexually recombine and increase their genotypic diversity much more frequently than is likely in dilute open ocean waters. Associated with these bloom events, density-related hazards of resource competition and selection for antagonist species heighten the value and need for rapid diversification of cell populations. Blooms represent not only a response to advantageous growth conditions but are hotspots for sexual reproductive potential and genetic exchange and thus play an outsized role in shaping the resilience and adaptability of these organisms.

## Acknowledgements

We are grateful to Mrunmayee Pathare for technical assistance with the Imaging FlowCytobot, and to Dr. Matthew Johnson for providing training and access to the ThermoFisher Attune Cytpix flow cytometer. This paper is a result of research funded by the National Oceanic and Atmospheric Administration National Centers for Coastal Ocean Science Competitive Research Program under award NA19NOS4780182 to the Virginia Institute of Marine Science. This is ECOHAB publication number ECO1125. Contributions by W.G.Z. were funded by the Qiushi Feiying project at Zhejiang University.

## Author contributions

S.S.C. and M.B. devised the study. S.S.C, N.A., and M.B. designed the experimental setup. M.B. collected field data. S.S.C, N.A., and W.G.Z. performed the experiment. S.S.C. processed samples and analyzed culture and field data. M.B. supervised the project. S.S.C. led manuscript writing. All authors provided input on the manuscript.

