## Supplementary Figures for "Rapid sexual reproduction in a mixotrophic dinoflagellate revealed through temporal partitioning of cellular processes"

### Supplementary Material

Serena Sung-Clarke[1,2], Nour Ayache[1], Wenguang Zhang[3], Mengmeng Tong[3], Juliette L. Smith[4], and Michael Brosnahan[1]

[1] Woods Hole Oceanographic Institution Biology Department, MA, USA, [2] MIT-WHOI Joint Program in Oceanography/Applied Ocean Science & Engineering, Cambridge and Woods Hole, MA, USA, [3] Zhejiang University Ocean College, Zhoushan, Zhejiang, China [4] Virginia Institute of Marine Science, William & Mary, Gloucester Point, VA, USA

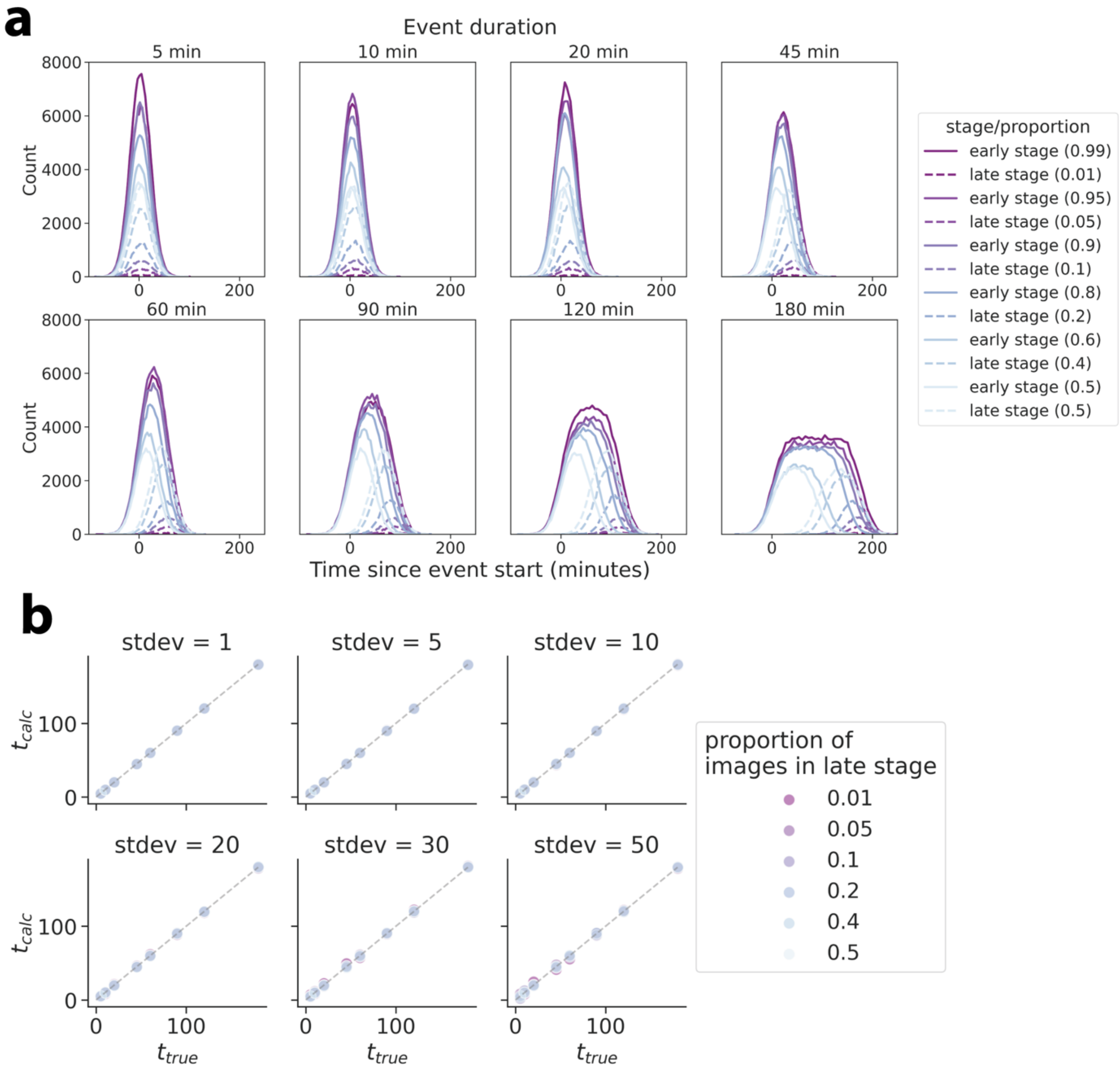


**Supplementary Figure 1.** Toy model simulation assessment of the method deriving an event duration from the difference in the midpoint between two known sub-stages of that event. **(a)** timing of predicted distributions of early and late stage observations across 8 different event durations and varying relative frequencies of early and late stage observations (sd = 20). **(b)** A calculated duration from the means of the early and late stage observation timing, where $t_{calc} = 2(\mu_{late} - \mu_{early})$, and standard deviations are varied. This method is robust across relative proportions in event stages and distribution variance.

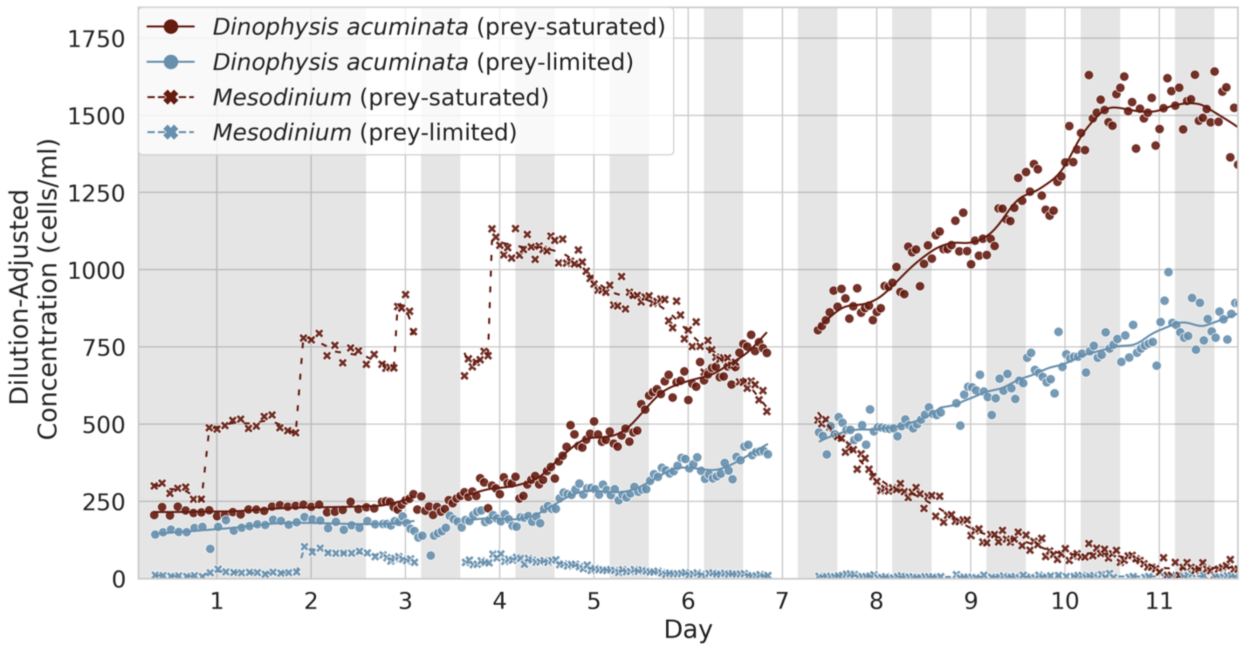


**Supplementary Figure 2.** Dilution-adjusted concentration of *Dinophysis* and *Mesodinium* in the prey-saturated and prey-limited treatments in the culture experiment, which was used to calculate accumulation and grazing rates.

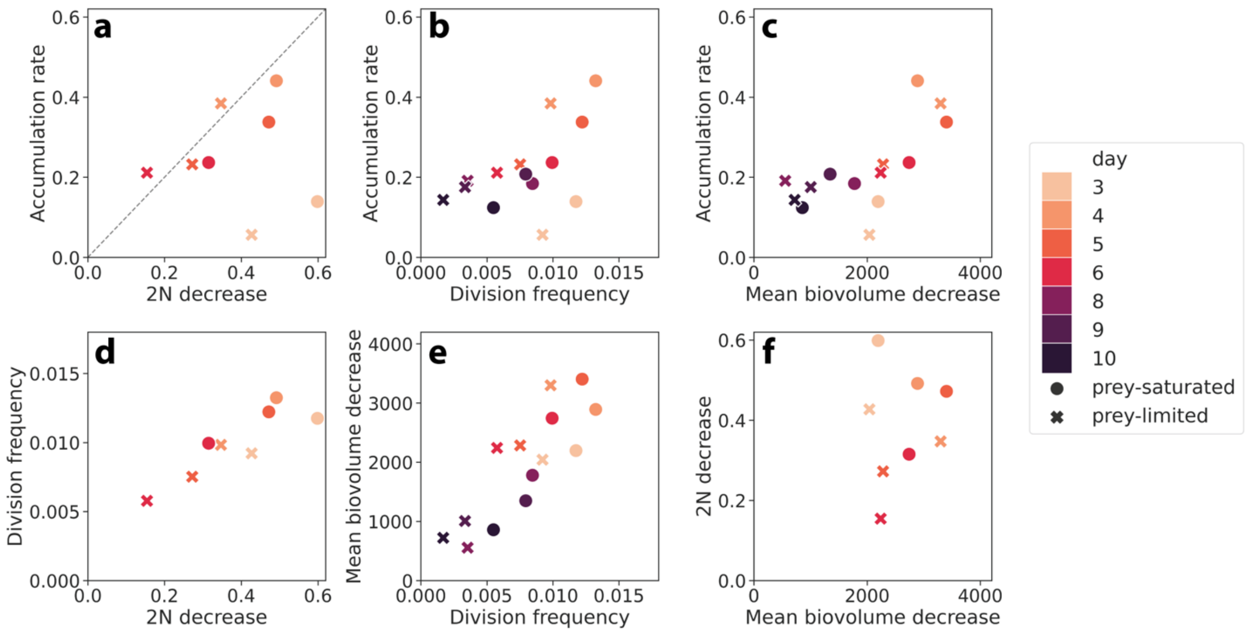


**Supplementary Figure 3.** Comparison of several metrics of growth estimates from the culture experiment, namely mean frequency of division, accumulation rate, daily biovolume decrease from maximum to minimum, and daily reduction in 2N proportion from maximum to minimum. Dashed line in the top-left plot is a 1:1 line.
